# Genetically encoded affinity reagents (GEARs): A toolkit for visualizing and manipulating endogenous protein function *in vivo*

**DOI:** 10.1101/2023.11.15.567075

**Authors:** Curtis W. Boswell, Caroline Hoppe, Alice Sherrard, Liyun Miao, Mina L. Kojima, Srikar Krishna, Damir Musaev, Ning Zhao, Timothy J. Stasevich, Antonio J. Giraldez

## Abstract

Probing endogenous protein localization and function in vivo remains challenging due to laborious gene targeting and monofunctional alleles. Here, we develop a multifunctional, universal, and adaptable toolkit based on genetically encoded affinity reagents (GEARs). GEARs use nanobodies and single chain variable fragments (scFv), which recognize small epitopes, enabling fluorescent visualization and selective degradation of protein targets in vivo. Furthermore, we delineate a CRISPR/Cas9-based epitope tagging pipeline to demonstrate its utility for producing knock-in alleles that have broad multifunctionality. We use GEARs to examine the native behaviour of the pioneer transcription factor Nanog and the planar cell polarity protein Vangl2 during early zebrafish development. Together, this toolkit provides a versatile system for probing and perturbing endogenous protein function while circumventing challenges associated with conventional gene targeting and is broadly available to the model organism community.

---

Understanding how proteins behave and function *in vivo* within their native, subcellular context is a continual challenge in model organism biology. While many animal systems are amenable to overexpression and transgenesis, these approaches can introduce artifacts due to sub- or supra-physiological expression levels^1–3^. Genome editing by site-specific nucleases has revolutionized genetics by permitting bespoke DNA editing abilities in a wide range of species^4–6^. These technologies enable the generation of tagged fusion proteins to study their endogenous function^7–9^. However, significant limitations such as laborious cloning and low germline transmission make precise gene editing by knock-in a challenge, especially when introducing larger fusion domains such as fluorescent proteins or degrons^10,11^. Although there have been expanded applications of green fluorescent protein (GFP) with targeted binding reagents such as degradation^12^, transcriptional activation^13^ and proximity interaction mapping^14^, most other tags such as DsRed, optogenetically-inducible domains^15^ and the auxin-inducible degron (AID)^16^ are monofunctional. This limited flexibility requires re-derivation of new alleles for each application. Thus, there is a significant need to improve reagents and methodologies to enable diverse experimental paradigms.

The aforementioned nanobody-based binding reagents developed for GFP represent an example of expanding the functionality of this tag beyond fluorescent imaging. Although GFP can be introduced into certain organisms more easily using genome editing^9^, pipelines for generating GFP alleles remains challenging and inefficient in systems such as zebrafish, medaka and mouse. Although the introduction of GFP into the genome is hindered by its size, shorter sequences can be introduced with greater ease and higher efficiency^17,18^. Hence, we sought to develop a methodology that allows researchers to create a multifunctional endogenous protein manipulation system with superior versatility and adaptability.

Here we develop a toolkit of **<u>g</u>**enetically **<u>e</u>**ncoded **<u>a</u>**ffinity **<u>r</u>**eagents, referred to as GEARs. This platform consists of short epitopes that recruit a wide variety of adapters such as fluorophores, degrons or HaloTags^19^ with high specificity and affinity through fusions to nanobodies or single chain variable fragments (scFvs). Furthermore, we establish a pipeline to efficiently knock-in epitope tags to genes of interest. By targeting genes using conventional CRISPR/Cas9 reagents and a single-stranded donor oligonucleotide (ssODN), we precisely engineer epitope-tagged alleles of *nanog* and *vangl2*. Using GEARs, we characterize the endogenous expression and localization of the encoded proteins and visualize Nanog dynamics during genome activation in zebrafish.

GEARs are broadly applicable to a wide range of animals and experimental designs and thus represent a powerful resource for the model organism and genetics community. This toolkit provides an easy plug-and-play approach allowing users to generate endogenously tagged alleles with limitless versatility. Together, GEARs-generated tagged alleles enable custom control over protein function *in vivo*, while circumventing many of the challenges that currently encumber conventional gene targeting.

## DESIGN

Historically, approaches to investigate endogenous protein function rely on antibody detection of epitopes 1) on the protein surface or 2) inserted as direct fusions to the locus of interest. Both approaches present with substantial limitations. Firstly, traditional antibody detection for microscopy is limited to fixed tissue and therefore can only provide static snapshots. Secondly, primary antibody availability can provide an experimental bottleneck, as some protein targets are less amendable to detection and producing new antibodies is laborious and expensive. Introducing a direct fusion tag such as GFP through genome engineering circumvents the need for protein specific antibodies because commercial anti-GFP antibodies are rapidly available. However, a large protein tag, such as GFP, increases the potential of interfering with the protein function and requires an in-frame knock-in of a long insert, tied to low insertion efficiency in some model organisms. The discovery of short epitope tags, such as HA^20^, together with developments of synthetically engineered small epitope tags, such as FLAG^21^ and ALFA^22^, overcome this challenge and allow for more efficient genome insertion while offering tags for a variety of applications. Together with the development of nanobodies and scFvs (single domain and single chain antibodies), short epitopes offer the potential to create truly versatile tags. Due to their size, they are less likely to affect the structure of tagged proteins, their localization, oligomerization, and protein-protein interactions.

Here we have characterized and multiplexed small epitope tags and their cognate tag-binding reagents, creating a multi-functional toolbox that can be applied to different model systems. Using codon-optimized epitope tags and their binders we first test their efficiency in zebrafish and mice. We further offer a direct comparison of tags and their efficiency in protein degradation *in vivo* and visualization in fixed and live cell experiments. Additionally, we developed a fully synthetic CRISPR/Cas9 approach to knock in these tags, increasing the methodologies to enable diverse experimental paradigms.

The aforementioned nanobody-based binding reagents developed for GFP represent an example of expanding the functionality of this tag beyond fluorescent imaging. Although GFP can be introduced into certain organisms more easily using genome editing^9^, pipelines for generating GFP alleles remains challenging and inefficient in systems such as zebrafish, medaka and mouse. Although the introduction of GFP into the genome is hindered by its size, shorter sequences can be introduced with greater ease and higher efficiency^17,18^. Hence, we sought to develop a methodology that allows researchers to create a multifunctional endogenous protein manipulation system with superior versatility and adaptability.

Here we develop a toolkit of **g**enetically **e**ncoded **a**ffinity **r**eagents, referred to as GEARs. This platform consists of short epitopes that recruit a wide variety of adapters such as fluorophores, degrons or HaloTags^19^ with high specificity and affinity through fusions to nanobodies or single chain variable fragments (scFvs). Furthermore, we establish a pipeline to efficiently knock-in epitope tags to genes of genome integration efficiency compared to traditional methods. Our modular system provides researchers with the opportunity to add novel tools in the future, including but not limited to immunoprecipitation, mass spectroscopy and super-resolution microscopy.

## RESULTS

### GEARs function in vivo to detect exogenously expressed protein targets

To establish whether GEARs would be functional and non-toxic to embryonic development, we surveyed a variety of single chain variable fragment (scFv) and nanobody (Nb) pairs (referred to as binders) with their respective peptide epitopes (Figure 1A). To this end, we tested whether the anti-HA scFv^23^ (referred to as FbHA), anti-FLAG scFv^24^ (referred to as FbFLAG), anti-ALFA Nb^22^ (referred to as NbALFA), anti-VHH05 Nb^25^ (referred to as NbVHH05) and anti-127d01 Nb^26^ (referred to as Nb127d01) binders would (1) fold correctly and localize homogeneously in the cell and (2) detect exogenous targets *in vivo* (Figure 1B). First, we generated codon-optimized binders fused to EGFP and added a stabilizing 3’UTR (see methods) to ensure robust and lasting expression for several days during embryogenesis without the need of a stable transgene. Next, we synthesized mRNA and injected 1-cell staged zebrafish embryos that were imaged at 6 hours post fertilization (hpf). We detect diffuse cytoplasmic and nuclear fluorescence for all constructs, suggesting that these GEARs are well tolerated, fold properly *in vivo* and are not excluded from subcellular compartments (Figure 1C-G). Notably, the FbHA appeared to form one to two distinct foci in close proximity to the nuclei as well as the mitotic apparatus (Figure S1A) which we attributed to off-target binding of centriolar proteins, a phenomenon observed with crossreactivity in some commercial antibodies^27,28^. Overall, expression of these binders did not reveal phenotypic effects at the injected concentrations. Together, these results demonstrate that these five binders can fold and function *in vivo* at physiological temperatures and pH different from their initial design (37***°***C compared to 28***°***C rearing temperature of zebrafish embryos).

**Figure 1:**
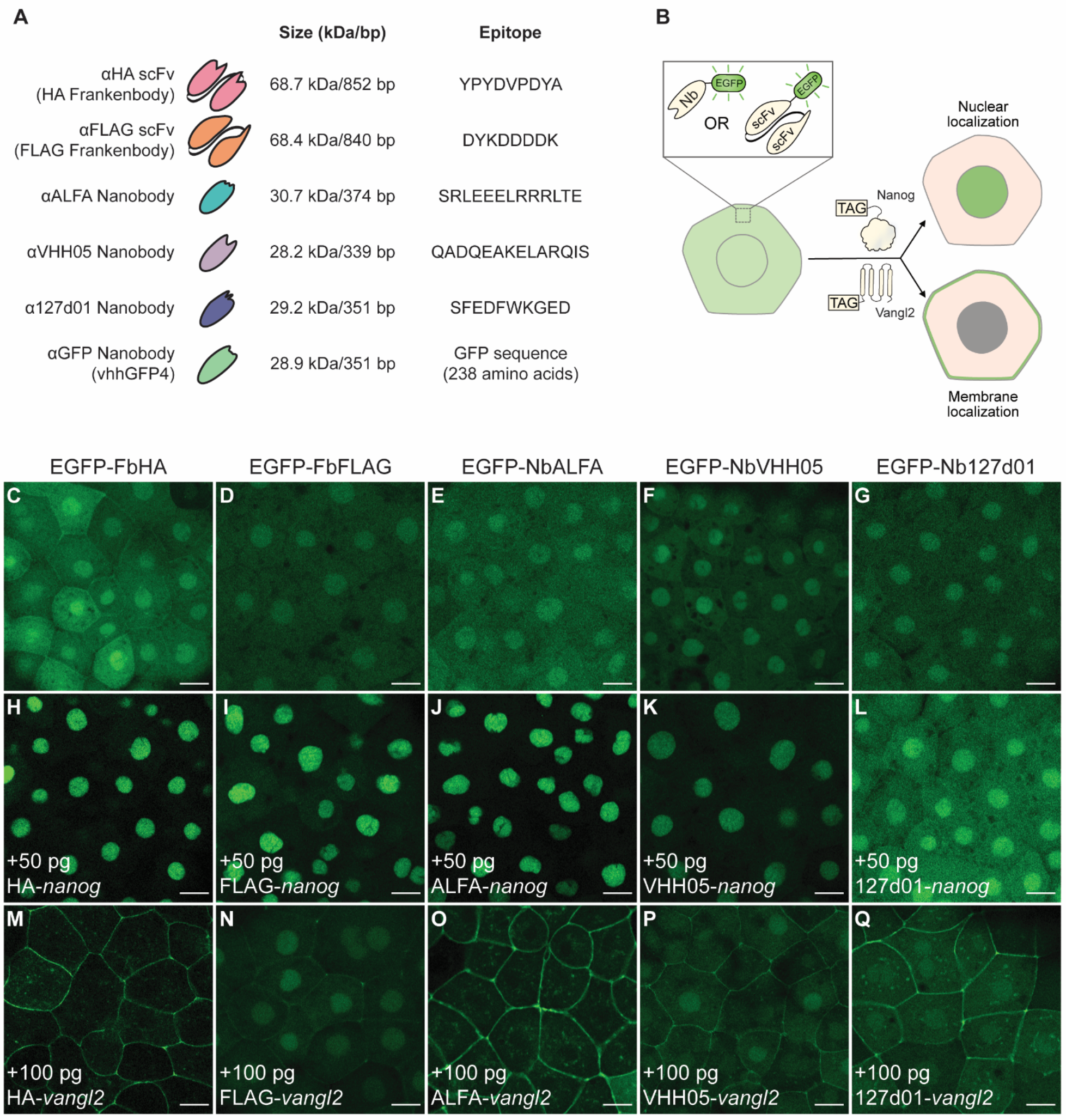
Genetically encoded affinity reagents (GEARs) function *in vivo*. (A) Overview of genetically encoded probes, their respective size and their target epitopes. (B) Schematic of assay for visualizing GEAR binding *in vivo*. Nanobodies or scFvs were fused to EGFP and injected into wildtype zebrafish embryos, either alone or with tagged versions of nuclear (Nanog) or membrane (Vangl2) targets. Localization of EGFP reflects the *in vivo* binding ability of the GEAR. (C-G) Localization of EGFP-GEAR ((C) FbHA, (D) FbFLAG, (E) NbALFA, (F) NbVHH05 and (G) Nb127d01) in wildtype embryos at 6 hpf without target introduction. (H-L) Localization of EGFP-GEAR ((H) FbHA, (I) FbFLAG, (J) NbALFA, (K) NbVHH05 and (L) Nb127d01) in wildtype embryos at 6 hpf co-injected with 50 pg *nanog* mRNA with cognate target epitope tags. (M-Q) Localization of EGFP-GEAR ((M) FbHA, (N) FbFLAG, (O) NbALFA, (P) NbVHH05 and (Q) Nb127d01) in wildtype embryos at 6 hpf co-injected with 100 pg *vangl2* mRNA with cognate target epitope tags. Scale bar, 20 □m (C-Q). See also Figure S1.

Next, we tested whether GEARs could bind to their cognate tags *in vivo* at various subcellular localizations. We cloned epitope tags for each of the GEARs onto the N-terminus of zebrafish *nanog* and *vangl2*. Nanog is a maternally deposited transcription factor that has pioneering activity in the embryo and regulates genome activation, localizing to the nucleus^29^. Vangl2, a core component of the planar cell polarity pathway, is localized to the membrane^30^. Importantly, the biological function of Nanog and Vangl2^30^ are unperturbed upon tagging the N-terminus of the protein (Figure S1 B-E). We reasoned that nuclear or membrane translocation of the EGFP-GEARs would provide a robust readout for binding *in vivo*. We co-injected the EGFP-tagged GEAR constructs into 1-cell staged embryos with their cognate tagged *nanog* mRNAs and found that all GEARs translocate to the nucleus with varying efficacy (ALFA/HA/FLAG>VHH05>127d01; Figure 1H-L). Furthermore, tagged *vangl2* mRNA also induced EGFP-tagged GEARs to translocate to the membrane (ALFA/HA/127d01>VHH05> FLAG; Figure 1M-Q). 127d01 and FLAG GEARs display varying efficiencies depending on the cellular localization of the epitope tagged protein. Overall, the ALFA Nb exhibited the most robust signal for both targets *in vivo* while displaying the least amount of background (Fig 1J,O). These results demonstrate the applicability of using GEARs to detect a variety of epitopes and targets during zebrafish development.

Next, we examined whether the nanobody-based GEARs would be compatible with fusions to other adapter proteins, to expand the gamut of fluorophores. To this end, we replaced the EGFP in the ALFA, VHH05 and 127d01 GEAR constructs with ORFs that encode mNeonGreen^31^, mScarlet-I^32^ and mTagBFP2^33^. When co-injected with tagged *nanog* mRNA, all fluorescent protein fusions were localized to the nucleus (Figure S1, F-H, J-L, N-P) suggesting that GEARs are amenable to multiple fluorescent adapters. Additionally, we generated HaloTag fusions for the nanobody-based GEAR binders. HaloTags can bind to a wide range of substrates such as immobilized surfaces for protein purification, reactive ligands for protein:protein interaction mapping and fluorescent dyes for super resolution imaging^19^. When co-injected into 1-cell staged embryos with or without tagged *nanog*, and incubated with the fluorescent JFX650 dye^34^ (which recognizes and bind the HaloTag) we observed robust fluorescent signal (Figure S1I, M, Q). These results suggest that GEARs are amenable to a wide variety of fluorescent adaptors and cargo proteins.

### GEARs can bind and degrade target proteins with high efficiency

Given the diverse cargo that could be bound by GEARs, we wondered if these binders would be amenable for targeted protein degradation. Recently, several genetic systems have adapted anti-GFP nanobody systems to engage targeted protein destruction in flies^12^, worms^35^, fish^36^ and human cells^37^. While these strategies rely on binding to GFP-tagged proteins, integration of large tags remains challenging in several model systems. Thus, it would be more effective to develop reagents that rely on shorter epitope tags that can be integrated with greater efficiency such as those offered by GEARs.

To test whether GEARs could facilitate degradation of tagged proteins, we adapted the zGrad GFP nanobody system^36^. zGrad uses the zebrafish F-box protein Fbxw11b fused to an anti-GFP Nb to target GFP-tagged proteins via ubiquitinylation for proteasomal degradation. Hence, we fused *fbxw11b* to GEAR binders. We titrated the expression of these mRNAs by injection into wild-type zebrafish embryos and found that all constructs are well tolerated without inducing ectopic phenotypes or toxicity (Figure S2A).

To test degradation, we employed a bicistronic reporter encoding membrane-tdTomato and GEAR epitope-tagged H2B-GFP split by a T2A self-cleaving peptide in zebrafish and mouse (Figure 2A-C). This reporter expresses 1:1 stoichiometric amounts of membrane and tagged nuclear fluorescent proteins (Figure 2D, I). Upon expression of a degrader, loss of nuclear GFP signal would indicate effective protein degradation. Upon co-injection of the bicistronic mRNA reporter with GEAR degrader mRNAs (referred to as ALFAgrad, VHH05grad and 127d01grad), we observe loss of GFP signal from the zebrafish embryos, with varying efficiency (Figure 2E-G). We observed 90%, 78% and 50% reduction in EGFP signal with ALFAgrad, VHH05grad and 127d01grad respectively at 10 hpf (Figure 2H and S2B-D), and this effect was maintained for over 24 hours (Figure S2G-I). The efficiency of GEARgrads (ALFAgrad, VHH05grad) was comparable to that of zGrad (Figure S2E-J). Because GEARgrad relies on the zebrafish F-box protein, it was unclear whether it can efficiently recruit ubiquitinylation machinery in other model systems. To address this, we injected the ALFA-tag reporter mRNA into one-cell stage mouse zygotes and the ALFAgrad in one of the two cells at the 2-cell stage (Figure 2C), such that the uninjected cell acts as no degradation control. Strikingly, we saw robust clearance of nuclear EGFP signal in the degron injected side (Figure 2I-I’’’), with a clearance efficiency of 96% (Figure 2J). These results demonstrate the flexibility of the GEARs to target proteins across different model systems, where ALFAgrad works most efficiently.

**Figure 2:**
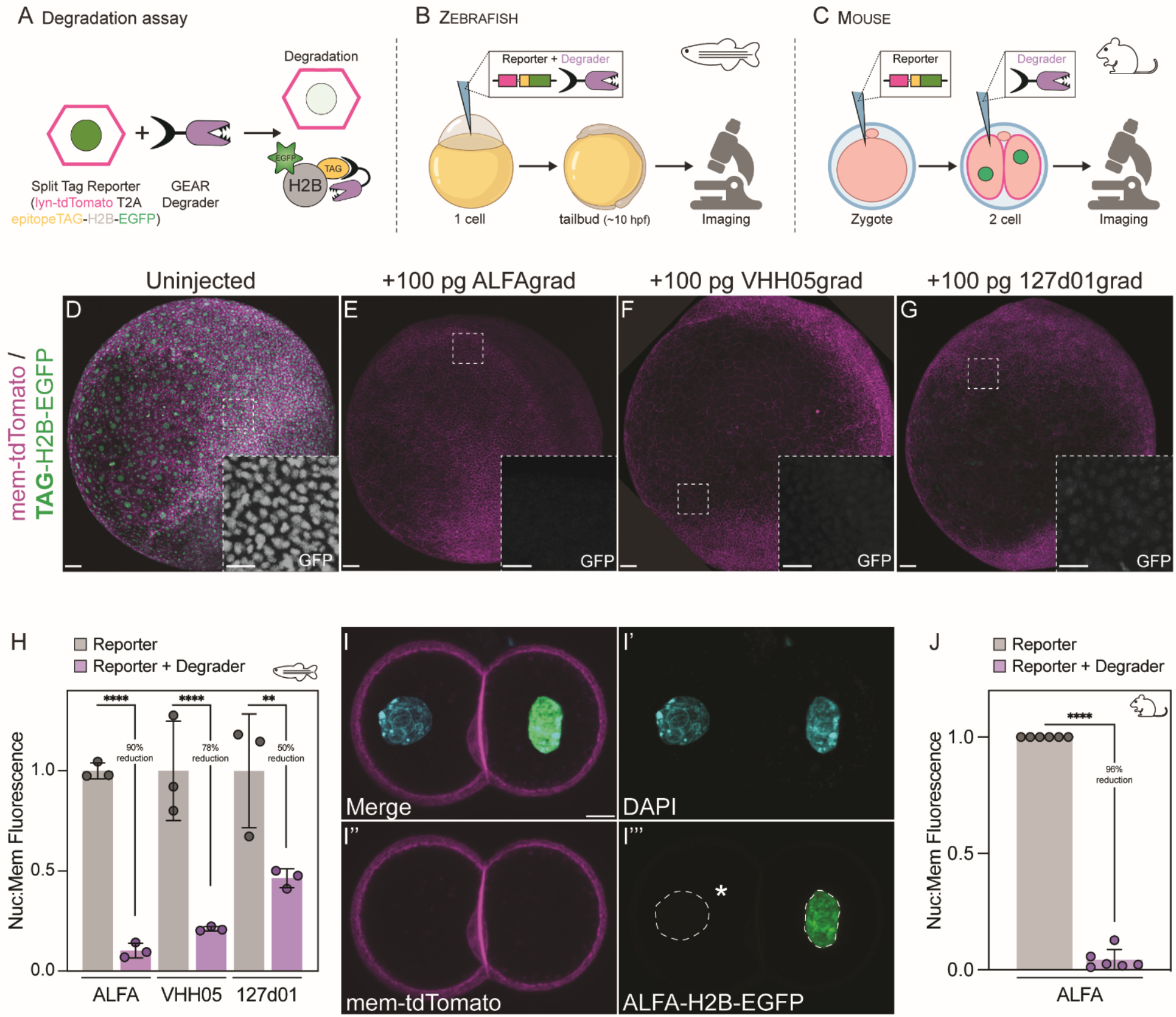
GEARs function in targeted protein depletion across vertebrate systems. (A) Schematic outlining the degron assay. A split reporter consisting of a membrane-targeted tdTomato (lyn-tdTomato) and an epitope-tagged nuclear EGFP (TAG-H2B-EGFP) are separated by a T2A peptide sequence, producing stoichiometric amounts of membrane and nuclear proteins. Upon introduction of a GEAR degrader, the epitope-tagged nuclear EGFP will be cleared and the ratios of tdTomato:EGFP reflect the efficiency of degradation. (B) Schematic outlining zebrafish degron assays. Embryos are injected at the 1-cell stage with 50 pg split reporters, with or without cognate GEAR degrader. Embryos are grown to 10 hpf and then imaged for total tdTomato and EGFP fluorescence. (C) Schematic outlining mouse degron assays. Embryos are injected at the 1-cell stage with 50 pg split reporters, and then re-injected at the 2-cell stage into 1 of the 2 cells (with the uninjected cell serving as a no-degradation control) with the degron. Embryos are fixed at the late 2-cell stage, stained with DAPI and imaged for total tdTomato and EGFP fluorescence. (D-G) Representative maximum intensity projections of zebrafish embryos at 10 hpf injected with (D) split reporters alone, (E) the ALFA degrader [ALFAgrad], (F) the VHH05 degrader [VHH05grad] and (G) the 127d01 degrader [127d01grad]. (H) Quantification of zebrafish split reporter ± degrader assays. Data were adjusted to a mean of 1 for comparison and pooled from n=3 embryos for each condition. For individual biological replicate data see Figure S2B-D. (I-I’’’) Representative images of late 2-cell mouse embryos injected with ALFA split reporter at the 1-cell stage and then re-injected into a single cell at the 2-cell stage with ALFA degrader (left cell). (J) Quantification of mouse split reporter ± degrader assays. Data from n= 6 embryos. Scale bar, 50 □m (D-G) and 25 □m (insets of D-G), 10 □m (I-I’’’). Mean ± SD and Student’s t-test (H,J), ****p < 0.0001. See also Figure S2.

## GEARs can be coupled with gene targeting to visualize and manipulate endogenous proteins

Given the size of the epitopes recognized by GEARS (<14 amino acids), we sought to develop a rapid endogenous tagging method to achieve efficient gene targeting. To achieve this, we employed a knock-in approach using recombinant Cas9, synthetic single guide RNA (sgRNA) and a single stranded oligodeoxynucleotide (ssODN) as a donor template (Figure 3A). This fully synthetic approach enables a cost- and time-efficient method for introducing short epitope tags into endogenous loci (detailed method in supplementary information). As a proof-of-principle, we used the zebrafish *nanog* and *vangl2* loci to introduce single copies of GEAR epitopes to identify the tags that would prove most effective in a native context. To this end, we generated 3 precise N-terminal knock-in alleles of *nanog* and *vangl2* respectively: ALFA, VHH05 and 127d01 tags (Figure 3B, S3A-B). We recovered precisely tagged *nanog* alleles with an efficiency of 20% (ALFA), 5% (VHH05) and 12% (127d01), and of tagged *vangl2* with an efficiency of 14% (ALFA), 12% (VHH05) and 18% (127d01). These efficiencies are notably higher than conventional homologous recombination efficiencies (0.3%-16%) especially with the lack of a secondary reporter such as a fluorescent marker^11,38^. Tags were confirmed to be integrated in-frame by Sanger sequencing, and the alleles were bred to homozygosity (Figure 3B, S3A-B). Of note, maternal homozygous knock-in embryos (derived from homozygous mothers) proceeded through gastrulation normally and were phenotypically indistinguishable from wildtype embryos at 24 hpf, suggesting the alleles are functional (data not shown). This rapid tagging method enables the utilization of a single tagged allele to be paired with the wide range of GEARs reagents for multifunctional analysis.

**Figure 3:**
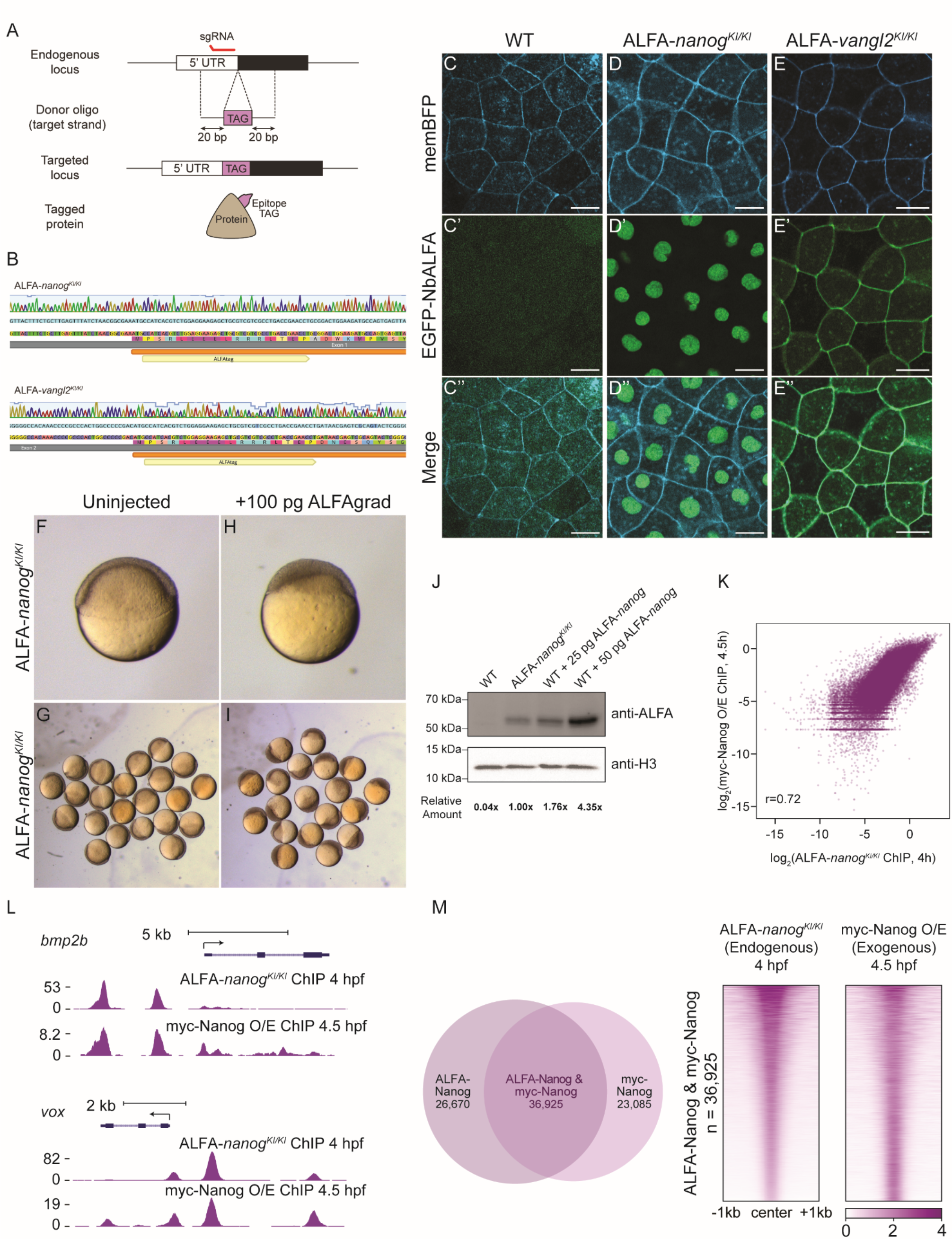
Genetically targeting endogenous *nanog* and *vangl2* with GEAR epitope tags produces versatile and multifunctional alleles. (A) Schematic outlining oligo targeting approach using single-stranded donor molecules and recombinant Cas9 to engineer tags into target genes. (B) Sanger traces of homozygous knock-in ALFA-*nanog* (top) and ALFA-*vangl2* (bottom) alleles. (C-C’’) Wildtype embryos at 8 hpf injected with 25 pg memBFP and 50 pg EGFP-NbALFA. (D-D’’) ALFA-*nanog*^*KI/KI*^ embryos at 8 hpf injected with 25 pg memBFP and 50 pg EGFP-NbALFA. (E-E’’) ALFA-*vangl2*^*KI/KI*^ embryos at 8 hpf injected with 25 pg memBFP and 50 pg EGFP-NbALFA. (F-G) Uninjected (F) individual and (G) group ALFA-*nanog*^*KI/KI*^ embryos at 6 hpf. (H-I) ALFAgrad injected (F) individual and (G) group ALFA-*nanog*^*KI/KI*^ embryos at 6 hpf. (J) Western blot comparing endogenously tagged Nanog to ALFA-Nanog reporters, injected at 25 pg and 50 pg concentrations. N= 25 embryos per condition. (K) Biplots showing correlations between ALFA-*nanog*^*KI/KI*^ ChIP-seq and Myc-Nanog overexpression (O/E) ChIP-seq. (L) Genome tracks of ALFA-*nanog*^*KI/KI*^ ChIP-seq and Myc-Nanog O/E ChIP-seq for Nanog target genes *bmp2b* (top) and *vox* (bottom). (M) Venn diagram shows the number and proportion of unique and shared Nanog ChIP-seq peaks in the genome. Heatmaps show ChIP-seq signal at shared Nanog peaks. Peaks were ranked based on the ALFA-*nanog*^*KI/KI*^ group. Scale bar, 20 □m (C-E). See also Figure S3 and S4.

We next focused our attention on the ALFA-tagged alleles, as they were shown to perform the best in exogenous experiments. First, using EGFP-tagged GEARs into embryos derived from homozygous knock-in mothers, we detected endogenous ALFA-*nanog* in the nucleus and ALFA-*vangl2* at the membrane in homozygous knock-in embryos (Figure 3C-E). These results demonstrate the ability for GEARs to target endogenous proteins *in vivo*. Next, we used ALFAgrad to test whether endogenously tagged *nanog* and *vangl2* alleles could recapitulate loss-of-function (LOF) mutant phenotypes. Injection of the ALFAgrad into ALFA*-nanog* homozygous embryos resulted in arrest of epiboly in 100% of the embryos phenocopying the maternal-zygotic *nanog* (MZ*nanog*) mutant phenotype (Figure 3F-I)^39,40^. Furthermore, ALFA-*vangl2* embryos injected with ALFAgrad fully recapitulated the MZ*vangl2* convergent-extension phenotype^30^ (Figure S3C-G). Importantly, the pool of maternal proteins is also targeted by these degron reagents, enabling robust clearance of proteins that establish the earliest events of embryogenesis. These data suggest that GEAR-mediated protein degradation can be as effective as generating a LOF mutant.

In zebrafish, Nanog has been investigated using exogenously provided reporter constructs due to a lack of primary antibody availability. However, using the anti-ALFA antibody (ALFA-Ab), we can now investigate endogenous Nanog directly. We used immunofluorescence (IF) and western blot to measure the amount of endogenous ALFA-tagged Nanog relative to exogenous Nanog used to rescue *nanog* mutant embryos at 4 hpf^29,40,44^. To our surprise, the endogenous concentration was nearly half the amount of exogenously provided *nanog* at 50% epiboly stage (5.3 hpf) (Figure 3J, S3H-M). Further, we performed a time-course analysis of Nanog protein and found that Nanog protein is present starting at the 1-cell stage and gradually increases over time, consistent with its role in genome activation^29^ (Figure S4A). These experiments demonstrate that ALFA-*nanog* knock-in fish report the physiological concentration of endogenously produced *nanog* protein, which is found at much lower levels than previously appreciated.

Nanog chromatin binding profile has been previously examined with ChIP-seq by overexpressing exogenous *nanog* in early zebrafish embryos^41^. Considering the different levels between endogenous and exogenous Nanog proteins, we asked whether the ALFA-Ab can be used to determine the DNA binding profile of endogenous Nanog using ChIP-seq and compared it to publicly available data using exogenous *nanog* expression. We observed correlation between these datasets (r=0.72) (Figure 3K) and a strong signal-to-noise ratio using the ALFA-Ab with 36,925 shared peaks representing ∼60% of the peaks in each sample (Figure 3L,M). Together these results suggest that the ALFA-Ab is a valuable tool for ChIP-seq.

### GEARs illuminate the native behaviour of Nanog during the earliest transcriptional events in zebrafish embryogenesis

Using the GEARS system, we set out to investigate the spatiotemporal dynamics of endogenous Nanog protein. Nanog is an ideal target to evaluate protein dynamics as (1) this pioneer transcription factor has intrinsically disordered regions that engage in concentration-dependent interactions and (2) exogenous fluorescently tagged Nanog localizes to subnuclear puncta during the maternal-to-zygotic transition^42,44^. While live imaging of exogenous Nanog has revealed important molecular behaviors, the localization of endogenous Nanog has not been studied due to the lack of appropriate tools.

We performed time-lapse imaging of endogenous ALFA-Nanog during the first three hours of development using the EGFP-NbALFA GEAR and compared it to exogenously expressed ALFA-Nanog or Nanog-mEmerald^42^ (Figure 4A, S4C).

**Figure 4:**
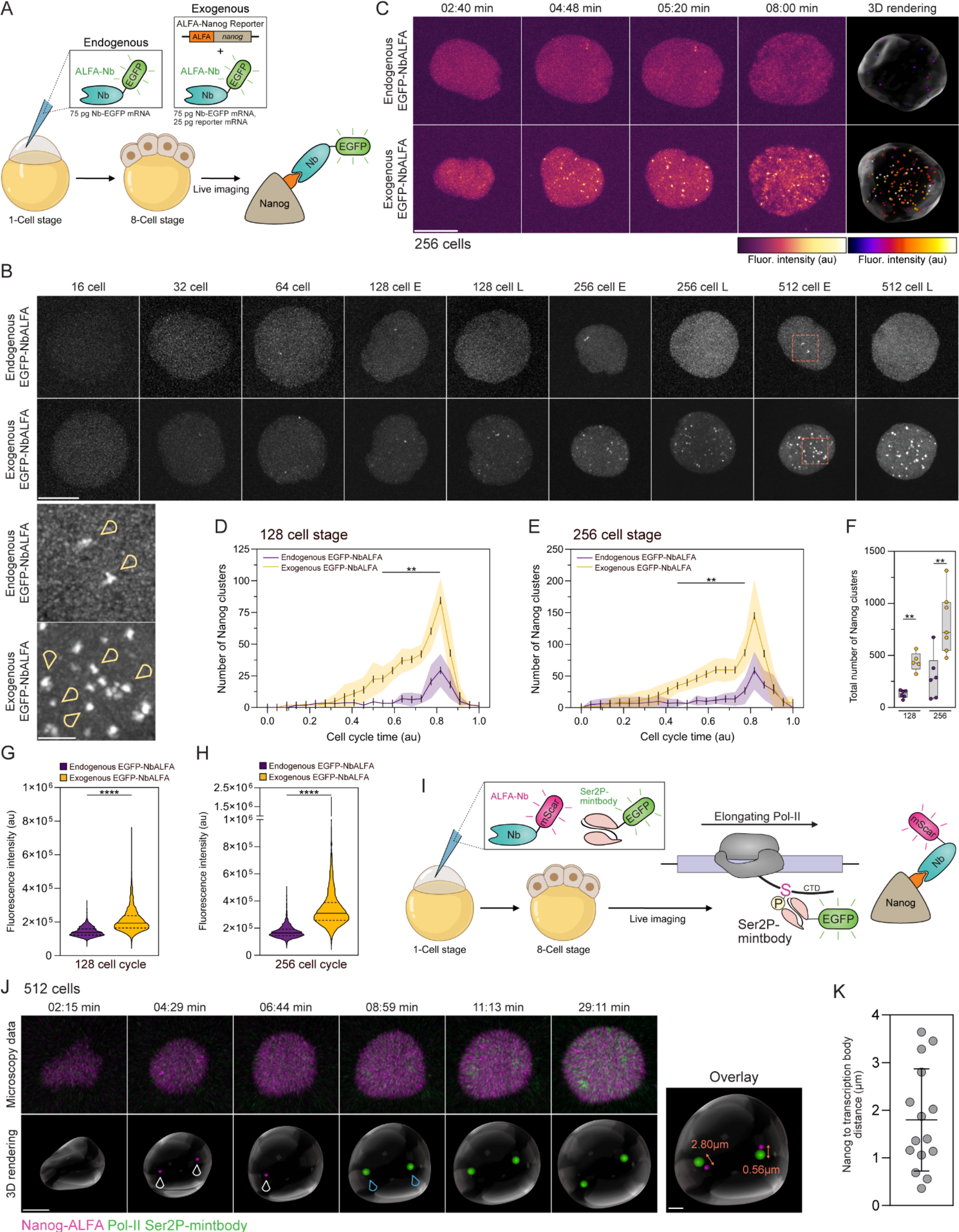
Endogenously tagged Nanog exhibits different behavior compared to overexpression. (A) Schematic outlining the live imaging setup. To visualize endogenous ALFA-Nanog, EGFP-NbALFA fusion protein mRNA was injected at the 1-cell stage into ALFA-*nanog*^*KI/KI*^ embryos. To visualize exogenous ALFA-Nanog, WT embryos were injected with an ALFA-*nanog* reporter (25 pg) and EGFP-NbALFA. All embryos were imaged continuously from the 8-cell to 1k cell stage. (B) Stills from live imaging datasets, comparing endogenous ALFA-Nanog and exogenous ALFA-Nanog. Images show comparable time points during different cell cycles (E=early, L=late; brightness and contrast were individually adjusted for better visibility and are therefore not comparable between stills. Insets shows zoomed areas in orange boxes and arrow heads indicate Nanog foci. (C) In detail comparison of time matched nuclei during 256-cell stage (stills from live imaging datasets) showing an increased number of Nanog foci in the overexpression condition compared to endogenously expressed ALFA-Nanog. A 3D rendering of the last time point is also shown. (D) Quantification of the number of visible Nanog foci during the 128-cell cell stage; n= 6 (endogenous) and 5 (exogenous) nuclei from 3 embryos. (E) As in (D) but for cell stage 256; n= 6 (endogenous) and 7 (exogenous) nuclei from 3 embryos. (F) Comparison of the total number of Nanog foci detected throughout cell cycles 128 and 256; n= 6 endogenous and 5 (128)/ 7 (256) exogenous nuclei from N=3 embryos. (G) Pooled fluorescence intensity data from all Nanog foci detected in cell cycle 128; n= 825 (endogenous) and 2235 (exogenous) Nanog foci from N=3 embryos. (H) As in (G) but for cell cycle 256; n= 2045 (endogenous) and 5885 (exogenous) Nanog foci from N=3 embryos. (I) Schematic of the live imaging setup visualizing endogenous Nanog in conjunction with elongating RNA Polymerase II (Pol-II). mScarlet-i3-NbALFA was injected into ALFA-*nanog*^*KI/KI*^ embryos at the 1-cell stage together with a mintbody recognizing Ser2P of the Pol-II C-terminal domain, fused to EGFP. (J) Stills of a representative nucleus during the 512-cell stage matched with a 3D rendering. Nanog foci (magenta, white arrow heads) precede transcriptional elongation of the *miR-430* locus (green, blue arrow heads). Nanog foci appear before the Pol-II signal but in very close 3D proximity (overlay of timepoints). In this example the 3D distances of Nanog foci to Pol-II transcription sites was 0.56 and 2.80 □m. (K) Quantification of data shown in (J); n= 15 Nanog/Pol-II measurements during the 512 or 1k cell stage from N=3 embryos. Bright Nanog foci are closely linked to the appearance of two large Pol-II transcription foci (mean=1.80 □m). Scale bar, 10 □m (B, C), 5 □m (J) and 2 □m (B and J inset). Mean ± SD (D,E,K), Median ± whiskers to min/max (F) and median ± interquartile range (G,H). **p < 0.01, ****p < 0.0001. Student’s t-tests (D,E) and Mann-Whitney test (F,G,H). See also Figure S4.

When providing Nanog exogenously (25 pg mRNA), we observed subnuclear foci formation starting at the 32-cell stage, forming characteristically bright foci at 128-512 cell stages (Figure 4B, S4D). Strikingly, endogenous Nanog accumulates in fewer foci starting at the 64-cell stage (Figure 4B) consistent with the lower concentration observed by IF and western blot. Interestingly, two bright fluorescent foci are robustly detected during early phases of cell stages 128-1k (Video1-3, Figure 4B, inset: yellow arrow heads). These likely represent the priming of *miR-430* transcription sites. The *miR-430* locus is the first zygotically transcribed region in the zebrafish genome and can be detected as early as the 64 cell cycle^43–45^.

Quantification of foci number over the cell cycle (128 and 256 cell stages) shows that endogenous Nanog protein forms significantly fewer foci. We observe a delay in the ramp up of foci formation, but it peaks at the same time relative to the exogenous expression, with an increase in foci number between cell cycles (Figure 4C-F, S4E-H, Video1-3). However, we cannot exclude that more micro foci are formed that are below the detection limit of this analysis. In addition to the stark increase in foci number, overexpression of Nanog also yielded an increase in foci fluorescence intensity, consistent with an overall increase of Nanog protein concentration (Figure 4G-H). While the injection of EGFP-nanobody increases the inherent background fluorescence due to unbound nanobody when compared to a direct fusion protein, we observe no significant differences in our two Nanog overexpression setups (Figure S4G-H), suggesting that the increased background of ALFA-nanobody does not obstruct the detection of Nanog foci. Together, these data suggest that overexpressed Nanog can either seed new foci or enlarge existing foci due to excess Nanog molecules.

### Endogenous Nanog foci prime the formation of large transcription bodies

In zebrafish embryos, elongating Pol-II is initially limited to two large, long-lived transcription bodies, which correspond to transcription of the *miR-430* gene cluster^43,44^. Exogenous Nanog transcription factor foci were shown to precede transcription body formation, consistent with its role as a pioneer transcription factor^29^. To investigate whether endogenous Nanog behaves identically, we used a fast-maturing mScarlet-i3-NbALFA GEAR^46^ to detect Nanog protein, as well as a genetically encoded Pol-II Ser2-EGFP reporter^42,47^ (Mintbody detecting RNA Pol-II phosphorylated on Serine 2) to visualize transcriptional elongation (Figure 4I). Like exogenous studies, we observed that Nanog foci precede transcription bodies in close proximity but are rapidly dissolved while transcription bodies are long lived (Figure 4J-K, Video4). These data support a model where *Nanog* binds to promoters and enhancers of *miR-430* and forms foci that are subsequently dissolved at the onset of transcription. Together, these results demonstrate the rapid mobility of *Nanog* protein during embryogenesis and illustrates the applicability of GEARs to visualize the behavior of endogenous proteins and gain biological insights.

## DISCUSSION

Here, we describe a toolkit and methodology that enables the generation of multifunctional and adaptable alleles. By surveying a variety of available genetically encoded binders, we identified those with the strongest performance *in vivo*. In addition to live imaging with these GEARS, we found that they are versatile in a variety of biological assays, such as by fusion to HaloTag or to degron adapters. Besides reagent development, we delineate a simplified pipeline for generating epitope-tagged alleles that can be directly used with GEARs. This framework for genetic engineering and probing is demonstrated by probing Vangl2 and Nanog during zebrafish embryogenesis. Together, these efforts enable a unified approach for biological interrogation of proteins in their native contexts.

Protein detection by epitope tagging has been frequently used for introducing short and inert sequences into genes. These tags are often small enough that they do not perturb function. For example, tagging of Wnt3 with a HA epitope allowed for the first visualization of an endogenous morphogen gradient which has been historically challenging to do with larger tags that disrupt Wnt3 function^48^. While these applications of epitope tagging can reveal static snapshots of biological phenomenon, advancement of technologies that detect these tags *in vivo* have lagged. Recent developments in these *in vivo* probing tools have been applied successfully in *Drosophila* and cultured cells^49,50^. These studies have applied similar principles of nanobody and scFv binding reagents to probe and manipulate proteins during development and homeostasis, and underscore the rapid acceleration of this field and the utility of these applications for a wide range of biological questions. For the first time in a vertebrate system, we have applied these *in vivo* probing tools with rapid knock-in editing to produce a multifunctional and versatile alleles.

A key parameter in protein targeting is ensuring that the resulting fusion retains biological function. We leveraged established tagging locations of *vangl2* and used loss-of-function rescue experiments with *nanog* mutants to determine the most optimal location for tag insertion. As S. Pyogenes Cas9 has sequence-specific requirements for DNA cleavage (the Protospacer Adjacent Motif (PAM) NGG), certain targets within the genome may be unavailable for targeting. Thus, additional Cas9 variants^51^ or additional nucleases (Cas12a)^52^ with differing cleavage sites may circumvent these sequence parameters and widen the targeting space. However, relative to homologous recombination (HR)-based methods, we observe a substantial increase in integration frequency (from an average of 0.3-16% to 10-20%) likely owing to the ease of integration of short DNA sequences into the genome through non-HR mechanisms. As zebrafish are less competent at engaging HR-dependent repair and favor microhomology-based repair during early embryogenesis^53,54^, oligo-based tagging likely occurs at much earlier stages of development and can increase the success of recovering germline transmitting events. Future optimizations of tagging locations, nuclease choice and germline screening methods will increase the throughput of allele generation. Together, this methodology enables the production of a single allele that can be used for a wide range of experimental paradigms.

In this study, we have introduced a novel degrader functionality into the epitope/nanobody field. This approach benefits from the small epitope tag size and strong affinity of nanobodies to their cognate tags. In both zebrafish and mouse embryos, ALFAgrad shows >90% efficiency in degrading nuclear proteins. GEARgrad provides several key advantages to other targeted protein depletion systems. First, in other degron systems such as dTAG^55^, HaloPROTAC^56^ and the AID system^16^, larger protein domains must be introduced into the gene of interest which can be time consuming and laborious. Second, some systems may require additional genetic components such as co-expression of accessory factors (like TiR1 with AID) often requiring depletion systems to be generated in transgenic backgrounds that express these components. Finally, these systems are often single-purpose and the established degron alleles cannot be repurposed for further experimental use beyond depletion. These features make established degron systems challenging to employ in model organisms where genome engineering is tied to low efficiency and emerging model systems where genome editing techniques are still being optimized.

Furthermore, we used GEARs to probe the behavior of endogenous Nanog protein. We observed two fluorescent Nanog foci during early embryonic cell cycles, which were detected in very close 3D proximity to two large transcription bodies, previously shown to correspond to transcription of the *miR-430* gene locus^43,44^. Interestingly, these two large Nanog foci, which most likely represent the priming of that gene locus for transcriptional activation, are the brightest foci observed during early embryogenesis, consistent with the high density of Nanog binding sites in this genomic region^29^. When providing a Nanog reporter, two foci from the larger pool of bright fluorescent foci were shown to precede *miR-430* transcription^42,44^. The increase in the number of Nanog protein foci compared to endogenous expression suggests that ectopic foci are formed due to increased concentrations of protein. One potential explanation is that small Nanog accumulations form at Nanog binding sites in the genome and excess unbound protein accumulates due to interactions of the proteins intrinsically disordered regions, growing these foci in size. Another potential explanation is that excess Nanog protein binds low affinity and ectopic sites in addition to canonical Nanog binding motifs. Previous work has shown that exogenous Nanog foci are DNA bound^42,44^ and therefore they are unlikely to represent protein accumulations in the nucleus. Given the recent increase in attention of the phase transition field to the formation of nuclear foci by transcription factors, it is worth considering that studying overexpressed proteins can lead to the formation of ectopic foci or the enlargement of existing foci due to excess molecules, highlighting the importance of studying endogenous proteins at their physiological concentrations, which can be achieved using the GEARs toolkit.

A major benefit to the GEARs methodology is the flexibility to introduce new components as they are released. Although we have highlighted this using multiple fluorescent and degron adapters, novel binders and cognate epitope can also be introduced into the system with ease. As the development of new intracellular probes continues, GEARs can be rapidly adapted to accommodate these components. Further, the development of chemically-inducible or light-inducible modification to the binding reagents would enable spatiotemporal control over GEAR activity. The breadth of options offered by GEARs ensures it remains nimble for future applications.

## LIMITATIONS

In this study, we have examined the use of GEARs for probing nuclear (Nanog) and membrane (Vangl2) protein targets. While these data demonstrate compelling *in vivo* binding to both compartments, it remains to be determined whether all other subcellular compartments are as amenable to engaging in a stable GEAR and target protein complex formation. It will therefore be useful to continue investigating additional targets with unique localization and determine if they are as amenable to binding GEAR components as shown here. Additionally, we demonstrate our epitope tagging approach on N-terminal targets. A thorough characterization of epitope tag location and GEAR efficacy will further our understanding of *in vivo* GEAR and target protein complex formation, and the effect of tag position on desired outcome.

## ACKNOWLEDGEMENTS

We thank Christopher Castaldi, Irina Tikhonova, and Bryan Szewczyk from the Yale Center for Genome Analysis for sequencing support and Sarah Dube, Timothy Gerson and Damilola Olowookere for zebrafish husbandry, and all members of the Giraldez lab for feedback and support. This project was supported by the Canadian Institutes of Health Research (CIHR) postdoctoral fellowship to C.W.B., the EMBO (ALTF 794-2021) and HFSP (LT0037/2022-L) postdoctoral fellowships to C.H., Surdna Foundation and Yale Genetics Venture Fund support for S.K., a National Institutes of Health grant R35 GM119728 to T.J.S., a National Institutes of Health grant R00 GM141453 to N.Z. and National Institutes of Health grant R01 HD100035 and National Institutes of Health grant R35 GM122580 to A.J.G.

## AUTHOR CONTRIBUTIONS

Conceptualization: C.W.B., C.H., A.J.G.; Formal analysis: C.W.B., C.H., L.M., M.K; Investigation: C.W.B., C.H., A.S., L.M., S.K., M.L.K.; Resources: D.M., N.Z., T.J.S.; Writing – Original Draft: C.W.B., C.H., A.J.G; Visualization: C.W.B., C.H., L.M., M.L.K.; Supervision: C.W.B., A.J.G.; Funding acquisition: A.J.G.

---

**Figure S1:**
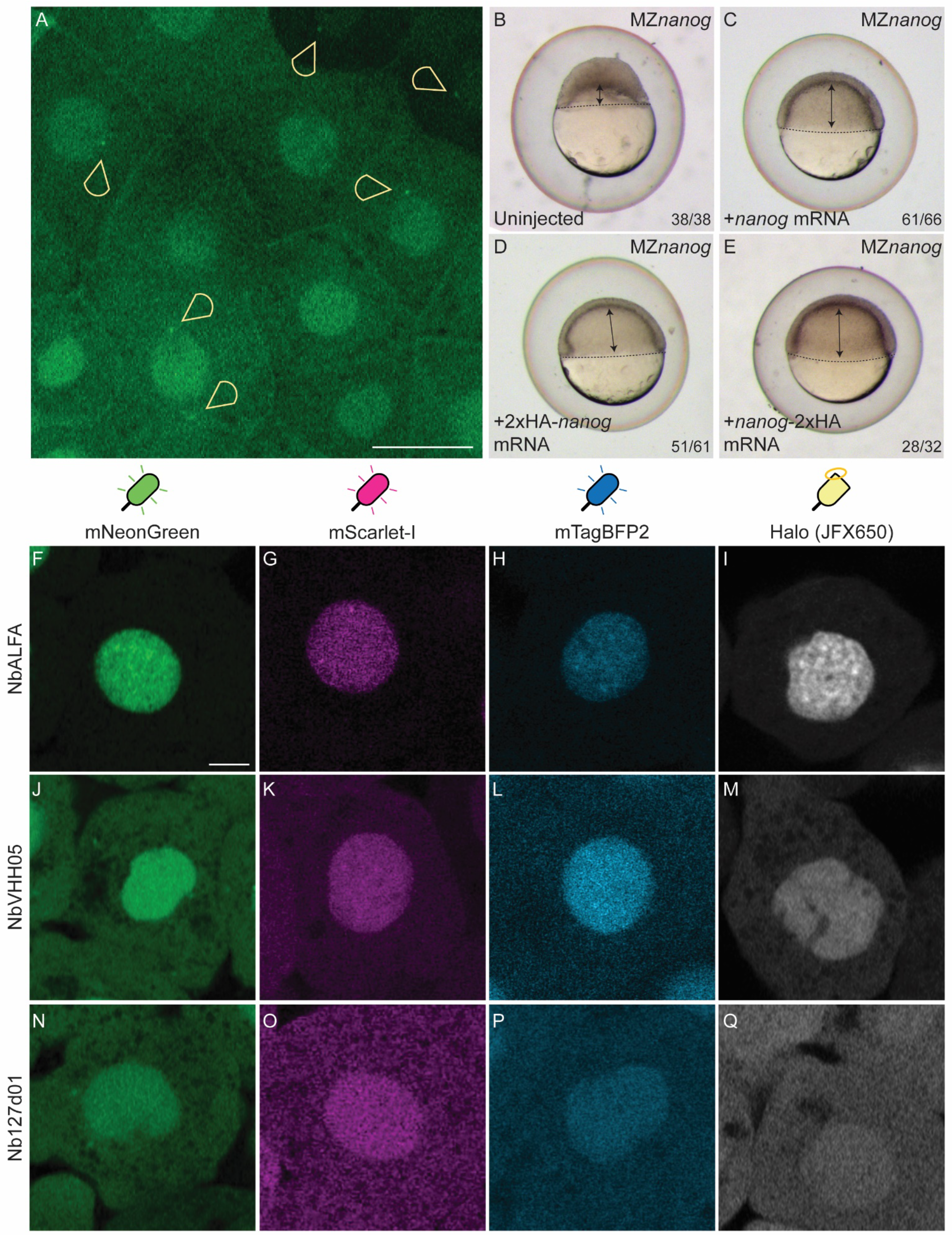
GEARs can be used with different adapters. Related to Figure 1. (A) EGFP-FbHA forms one to two distinct foci in close proximity to nuclei as well as the mitotic apparatus (yellow arrowheads). (B-E) The biological function of N-terminal and C-terminal tagged Nanog was compared to uninjected MZ*nanog* mutant embryos (B) that arrest before gastrulation. The mutant phenotype was rescued by injecting *nanog* mRNA (C), 2xHA tagged *nanog*-mRNA (D) and *nanog*-2xHA tagged mRNA (E). N numbers are indicated in the figure panels. (F-I) The ALFA GEAR adapter was fused to mNeonGreen (F), mScarlet-I (G), mTagBFP2 (H) or a Halo Tag (I) and injected together with the tagged *nanog* mRNA. (J-M) same as (F-I) but with the VHH05 GEAR adapter. (N-Q) same as (F-I) but with the 127d01 GEAR adapter. Scale bar, 20 □m (A) and 10 □m (F-Q).

**Figure S2:**
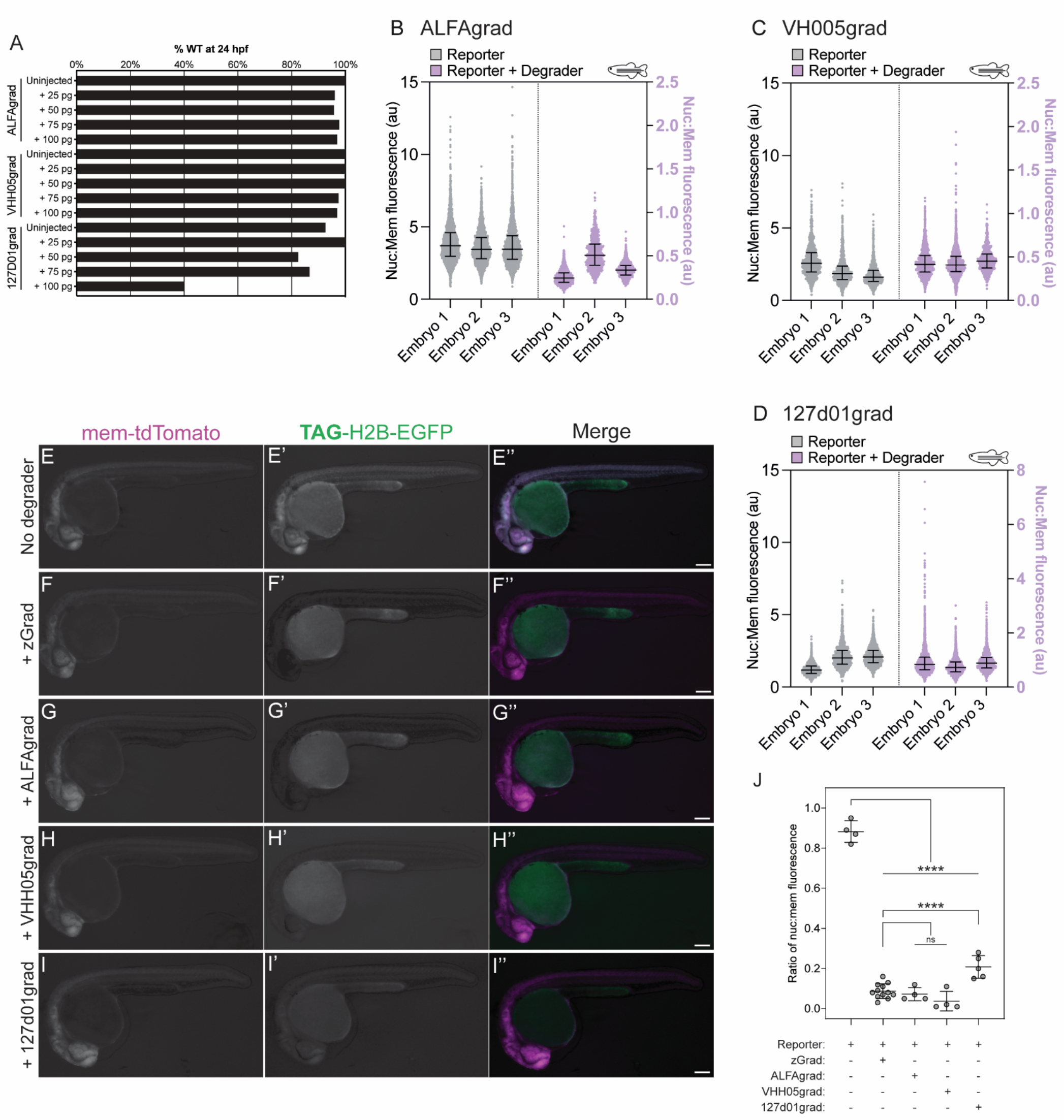
GEARgrad systems deplete nuclear proteins. Related to Figure 2. (A) Titration experiment to determine whether GEARgrad constructs are tolerated well *in vivo*. mRNAs were injected at different concentrations into WT zebrafish embryos and ectopic phenotypes were scored at 24 hpf. Data were collected from n=4 (uninjected), n=12 (zGrad), n=4 (ALFAgrad), n= 4 (VHH05grad) and n=5 (127d01grad) embryos. (B) Individual values of zebrafish split reporter ± degrader assay. The nuclear to membrane fluorescence ratio of individual nuclei is shown in reporter only injected embryos (R) and reporter + degrader injected embryos (R+D). Data shown from n= 2841, 2727, 2761 (R) and 1444, 662, 1523 (R+D) nuclei in 3 embryos respectively. Pooled data shown in Figure 2H. (C) As in B but for VH005grad with data from n= 946, 1007, 1050 (R) and 1377, 1862, 932 (R+D) nuclei in 3 embryos respectively. Pooled data shown in Figure 2H. (D) As in B but for 127d01grad with data from n= 1843, 1310, 1969 (R) and 2262, 1991, 1538 (R+D) nuclei in 3 embryos respectively. Pooled data shown in Figure 2H. (E-I) Representative maximum intensity projections of zebrafish embryos at 24 hpf injected with (E) split reporters alone, (F) GFP degrader [zGrad] (G) the ALFA degrader [ALFAgrad], (H) the VHH05 degrader [VHH05grad] and (I) the 127d01 degrader [127d01grad]. (J) Quantification of degron experiments (E-I) at 24 hpf. The ratio of nuclear to membrane fluorescence indicates the amount of nuclear protein degradation using different degrons. Data from n= 4 (CTRL), 12 (zGrad), 4 (ALFAgrad), 4 (VHH05grad) and 5 (127d01grad) embryos. Scale bar, 200 □m (E-I). Box with Tukey whiskers (box shows median, 25th and 75th percentile, + indicates mean; L) and mean ± SD (J). One-way ANOVA with a Dunnett’s multiple comparisons test shows the difference to the control or zGrad (J), ****p < 0.0001; ns, not significant.

**Figure S3:**
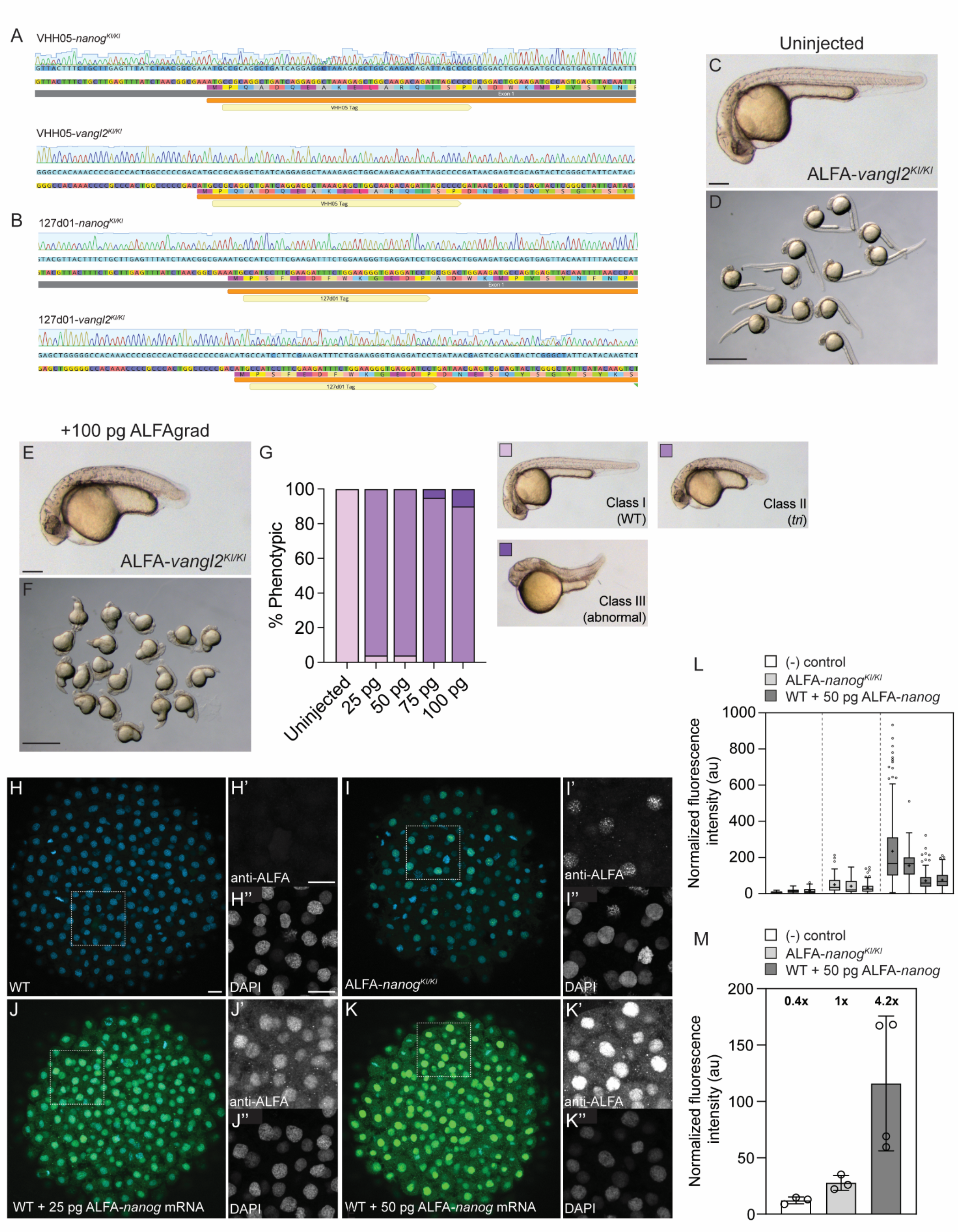
Tagging of *nanog* and *vangl2* with GEAR reagents permits the investigation of endogenous protein. Related to Figure 3. (A) Sanger sequencing traces of homozygous knock-in VHH05-*nanog* (top) and VHH05-*vangl2* (bottom) alleles. (B) as in (A) but for 127d01-*nanog* (top) and 127d01-*vangl2* (bottom) alleles. (C-D) Uninjected (C) individual and (D) group ALFA-*vangl2*^*KI/KI*^ embryos at 24 hpf. (E-F) ALFAgrad injected (E) individual and (F) group ALFA-*vangl2*^*KI/KI*^ embryos at 24 hpf. (G) Quantification of phenotypes observed with ALFA-Vangl2 depletion using ALFAgrad, in n=84 (uninjected), n=62 (25 pg), n=69 (50 pg), n=61 (75 pg) and n=48 (100 pg) embryos. (H-K) Representative IF images visualizing ALFA-Nanog (green) via an anti-ALFA Ab in uninjected WT (H), ALFA-*nanog*^*KI/KI*^ (I), 25pg ALFA-*nanog* mRNA injected (J) and 50pg ALFA-*nanog* mRNA injected (K) embryos. Nuclei stained with DAPI (blue). (L) Quantification of nuclear ALFA-Nanog signal in IF images of individual embryos. n= 145, 143, 191 (WT), 108, 77, 131 (ALFA-*nanog*^*KI/KI*^) and 167, 143, 125, 163 (50pg injection) nuclei from N= 3 (WT and ALFA-*nanog*^*KI/KI*^) or 4 (50pg injection) biological replicates. (M) Replicate data was pooled to compare endogenous ALFA-tagged Nanog relative to exogenously provided Nanog. Relative amount of ALFA-Nanog indicated in the graph and data from n= 3 (WT and ALFA-*nanog*^*KI/KI*^) or 4 (50pg injection) biological replicates. Scale bar, 200 □m (C,E), 1 mm (D,F), 100 □m (H-K) and 5 □m (H-K, inserts). Box with Tukey whiskers (box shows median, 25th and 75th percentile, + indicates mean; L) and mean ± SD (M).

**Figure S4:**
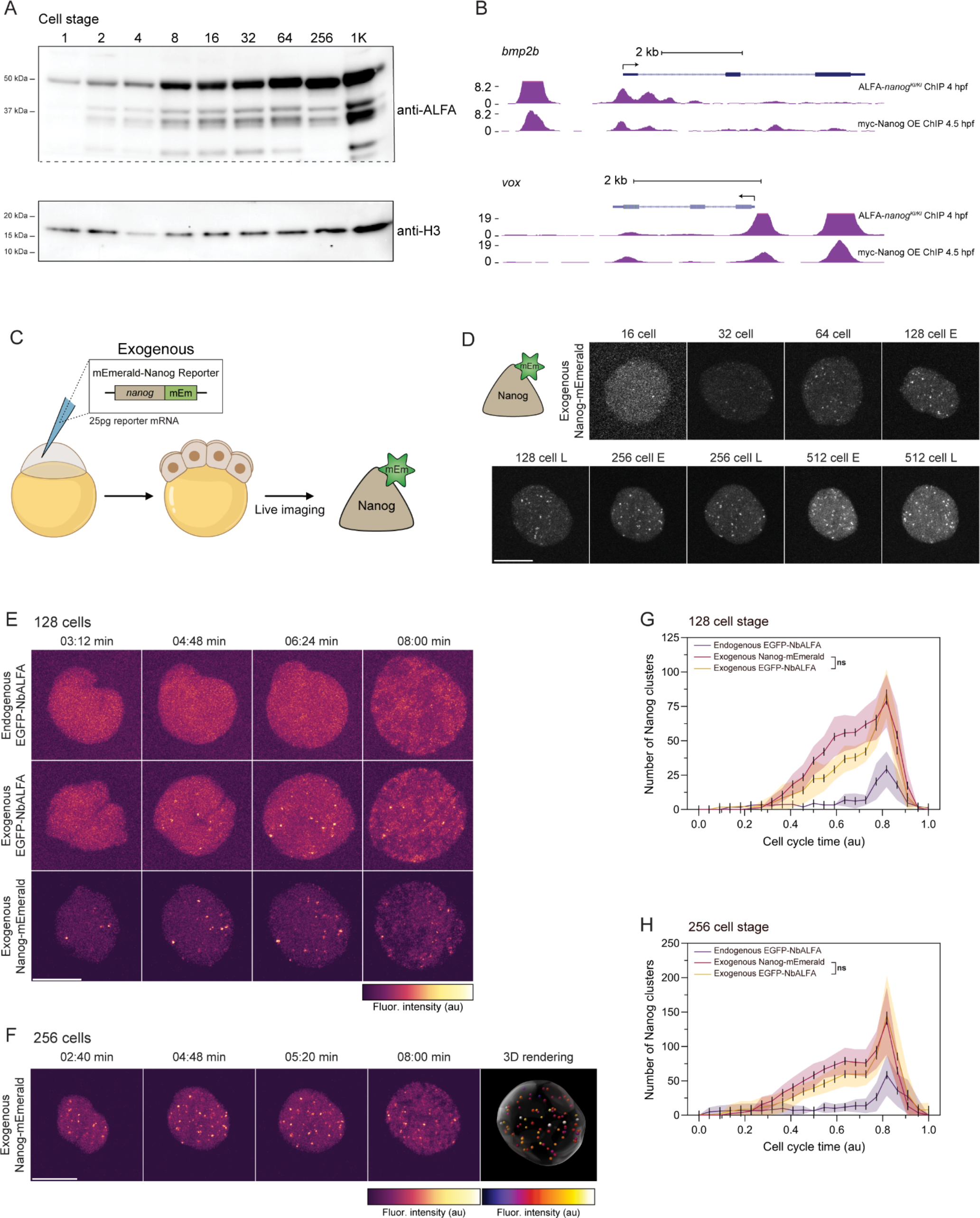
Investigating endogenous Nanog behaviour by western blot, ChIP-seq and live imaging. Related to Figure 3 and 4. (A)Western blot time course of endogenous Nanog concentrations detected with the anti-ALFA Ab. Each sample contains 15 embryos and H3 histone levels were used as the loading control. The membrane was cut at ∼25kDa (dotted line) prior to Ab incubation. (B) ChIP-seq genome tracks of Nanog target genes *bmp2b* (top) and *vox* (bottom), replicated from Figure 3L but at identical scales between conditions. (C) Live imaging of *nanog* overexpression. Embryos at the 1-cell stage were injected with mRNA encoding a direct Nanog and mEmerald fusion protein. Injected embryos were mounted at the 4-cell stage and imaged starting at the 8-cell stage. (D) Stills showing Nanog foci formation throughout cell cycles (E= early, L=late; brightness and contrast were individually adjusted for visibility and are not comparable between stills). (E) Time lapse still images comparing endogenous ALFA-Nanog, exogenous ALFA-Nanog and exogenous Nanog-mEmerald during cell stage 128. (F) Time lapse still images and a 3D rendering of Nanog-mEmerald injected embryos during cell stage 256. (G) Quantification of Nanog foci number over time during the 128 cell stage. Graph partially replicated from Fig 4D. (H) As in (G) but for Nanog foci number during the 256 cell stage. Graph partially replicated from Fig 4E. Scale bar, 10 □m (D-F). Mean ± SD (G,H). Student’s t-tests (G,H); ns, not significant.

## METHODS

### Lead Contact

Further information and requests for resources and reagents should be directed to and will be fulfilled by the Lead Contact, Curtis W. Boswell.

### Materials Availability

Plasmids and zebrafish lines generated in this study are available from the Lead Contact on request. Plasmid sequences will be included in the revised manuscript.

### Data and Code Availability

Raw ChIP sequence reads generated in this study will be made publicly available. Other materials generated in this study are available upon request.

## Experimental Model and Subject Details

### Zebrafish husbandry and maintenance

*Danio rerio* (zebrafish) embryos were obtained from natural matings of adult fish of mixed wild-type backgrounds (TU-AB and TLF strains) of mixed ages, ranging from 5-18 months. Zebrafish were maintained and used in accordance with the Yale University AAALAC guidelines under a protocol approved by the Yale University IACUC (protocol number 2021-11109). Embryos were grown and staged according to published standards^58^ and all zebrafish and embryo experiments were performed at 28°C.

### Mouse husbandry and maintenance

Mouse experiments were performed in compliance with ethical protocols approved by the Yale University Institutional Animal Care and Use Committee (IACUC) under protocol #2023-20324. Mouse embryos were generated by inducing hyperovulation in 4-week-old B6D2F1 females, which were mated with 8-week to 6-month-old B6D2F1 males. To induce hyperovulation, 5 IU of pregnant mare serum gonadotropin (PMSG) was injected intraperitoneally, followed by 7.5 IU of human chorionic gonadotropin (hCG) 47 h later. Zygotes were collected 20 h post-hCG and cumulus cells were removed using hyaluronidase diluted in M2 (Sigma, MR-051, Sigma, MR-015-D). Embryos were cultured in 25 μL drops of KSOM (MR-106-D, Sigma) covered with cell-culture grade paraffin oil (Copper Surgical, ART-4008-5P) in a cell culture incubator maintained at 37°C and 5% CO_2_.

## Method Details

### Molecular cloning

#### GEAR binders

DNA fragments containing the ORFs for FbHA, FbFLAG, NbALFA, Nb127d01 and NbVHH05 were codon-optimized using iCodon^57^ and purchased from IDT as GBlocks. Fragments were cloned into a custom pCS2+ EGFP expression vector as C-terminal fusions (EGFP-GEAR binder) using InFusion enzyme (Takara) and sequence verified. All reporters had an optimized *prrg2* zebrafish 3’UTR, selected from a pool of endogenous 3’UTRs (300-500bp length) that lacked known decay motifs (including *miR-430* target sequences (GCACT), AU-rich elements (ATTTA) and C-rich decay (CTCC, CCTC, CTGA, CACA, TCTC, ACTC, CTCT, CCTG) motifs). Unpublished data showed this 3’UTR increases mRNA stability and protein production several-fold compared to beta globin 3’UTR.For mRNA production, vectors were linearized with NotI and *in vitro* transcribed using mMessage mMachine SP6 kit (Ambion).

#### Fluorescent GEAR adapters

DNA fragments coding for mNeonGreen (Addgene plasmid # 128144), mScarlet-I (Addgene plasmid # 85044), mTagBFP2 (Addgene plasmid # 55295) and HaloTag (Addgene plasmid # 128603) were amplified, cloned into pCS2+ GEAR binder clones to replace EGFP ORF using InFusion enzyme (Takara) and sequence verified. For mRNA production, vectors were linearized with NotI (NEB) and *in vitro* transcribed using mMessage mMachine SP6 kit (Ambion). A DNA fragment for mScarlet-i3^46^ was codon-optimized using iCodon^57^, purchased from IDT as GBlocks and cloned into pCS2+ NbALFA as described above. pCS2+mNeonGreen-C Cloning Vector was a gift from Ken-Ichi Takemaru (http://n2t.net/addgene:128144; RRID:Addgene_128144). pmScarlet-i_C1 was a gift from Dorus Gadella (http://n2t.net/addgene:85044; RRID:Addgene_85044). mTagBFP2-Farnesyl-5 was a gift from Michael Davidson (http://n2t.net/addgene:55295; RRID:Addgene_55295). Nb-gp41-Halo (MoonTag-Nb-Halo) was a gift from Marvin Tanenbaum (http://n2t.net/addgene:128603; RRID:Addgene_128603)

#### Degron GEAR adapters and degron reporters

Zebrafish *fbxw11b* ORF was amplified from pCS2+ zGrad (Addgene plasmid # 119716) and cloned into pCS2+ GEAR binder clones to replace EGFP ORF using InFusion enzyme (Takara). Split fluorescent reporters were cloned by introducing ALFA, VHH05 and 127d01 epitope tags in-frame to the H2B-EGFP ORF in pCS2+TAG (Addgene plasmid # 26772). Vectors were sequence verified, linearized with NotI (NEB) and *in vitro* transcribed using mMessage mMachine SP6 kit (Ambion) for mRNA injection. pCS2(+)-zGrad was a gift from Holger Knaut (http://n2t.net/addgene:119716; RRID:Addgene_119716). pCS2-TAG was a gift from Shankar Srinivas; http://n2t.net/addgene:26772; RRID:Addgene_26772)

#### Nanog and Vangl2 fusions

Tandem HA tags were cloned as N-terminal or C-terminal fusions using InFusion enzyme (Takara) into a pCS2+ *nanog* expression vector^29^. Single HA, FLAG, ALFA, VHH05 and 127d01 epitope tags were cloned as N-terminal fusions using InFusion enzyme (Takara) into a pCS2+ *nanog* and pCS2+ *vangl2* expression vector. Vectors was sequence verified, linearized with NotI (NEB) and *in vitro* transcribed using mMessage mMachine SP6 kit (Ambion) for mRNA injection.

#### Live imaging reagents

Expression vectors containing *nanog*-mEmerald and the Pol-II PhosphoSer2 Mintbody-EGFP^42^ were linearized using NotI (NEB) and *in vitro* transcribed using mMessage mMachine SP6 kit (Ambion) for mRNA injection.

### GEAR localization experiments

#### Injections

Wildtype 1-cell staged embryos were dechorionated in pronase and injected with 50 pg of EGFP-GEAR mRNA alone or with 50 pg of tagged *nanog* mRNA or 100 pg tagged *vangl2* mRNA. Embryos were incubated at 28°C until they reached 30% epiboly (∼5 hpf) and mounted in 0.8% low melt agarose (GPG/LMP AmericanBio, AB00981-00050) in system water against a no. 1.5 cover slip.

#### Imaging

Embryos were imaged on a temperature controlled (28°C) upright Zeiss LSM 980 confocal microscope with an Airyscan 2 detector and a Plan-Apochromat 20x/0.8 M2 objective with bidirectional line scanning at a format of 4084 × 4084 pixels and 1.4x optical zoom. All images were collected at 16 Bit, and optical stacks were acquired at 0.5 μm spacing. EGFP fluorophores were excited using the 488 laser line at 4% laser power. Raw images were deconvolved using the Airyscan software. Individual representative slices of the enveloping layer are shown in Figure 1.

### GEAR multicolour adapter experiments

#### Injections

Wildtype 1-cell staged embryos were dechorionated in pronase and injected with 50 pg of tagged *nanog* mRNA alone Nov. 15, 23 or with 50 pg of mNeonGreen-GEAR, 50 pg of mScarlet-I-GEAR, 50 pg of mTagBFP2-GEAR or 50 pg of HaloTag-GEAR mRNA. Embryos were incubated at 28°C until they reached 30% epiboly (∼5 hpf) and mounted in 0.8% low melt agarose (GPG/LMP AmericanBio, AB00981-00050) in system water against a no. 1.5 cover slip. For HaloTag imaging, JFX650 (Janelia Fluor®) ligand was directly added at a concentration of 10 nM to low melt agarose and mixed thoroughly prior to mounting.

#### Imaging

Embryos were imaged on a temperature controlled (28°C) upright Zeiss LSM 980 confocal microscope with an Airyscan 2 detector and a Plan-Apochromat 20x/0.8 M2 objective with bidirectional line scanning at a format of 4084 x 4084 pixels and 1.4x optical zoom. All images were collected at 16 Bit, and optical stacks were acquired at 0.35 μm spacing. Fluorophores were excited using the 408 laser line (mTagBFP2) at 2% laser power, 488 laser line (mNeonGreen) at 1% laser power, 561 laser line (mScarlet-I) at 8% laser power and 639 laser line (Halo/JFX650) at 0.6% laser power. Raw images were deconvolved using the Airyscan software. Individual representative slices of the enveloping layer are shown in Figure S1.

### Degron experiments

#### Zebrafish injections

Wildtype 1-cell staged embryos were dechorionated in pronase and injected with 50 pg of ALFA, VHH05 or 127d01 split florescent reporter mRNA alone or with 100 pg of Fbxw11b-GEAR degron mRNA. For 10 hpf imaging, embryos were incubated at 28°C until they reached tailbud stage and mounted in 0.8% low melt agarose (GPG/LMP AmericanBio, AB00981-00050) in system water against a no. 1.5 cover slip. For 24 hpf imaging, embryos were incubated at 28°C until they reached Prim-6 stage and mounted in 3% methylcellulose on a glass depression slide.

#### Zebrafish imaging

For 10 hpf imaging, embryos were imaged on a temperature controlled (28°C) upright Zeiss LSM 980 confocal microscope with an Airyscan 2 detector and an EC Plan-Neofluart 10x/0.3 M27 objective with bidirectional line scanning at a format of 4084 x 4084 pixels and 1.0x optical zoom. All images were collected at 16 Bit, and optical stacks were acquired at 3.08 μm spacing. Fluorophores were excited using the 488 laser line (EGFP) at 5% laser power and 561 laser line (tdTomato) at 8% laser power. Raw images were deconvolved using the Airyscan software. Representative orthogonal projections of individual embryos are shown in Figure 2.

For 24 hpf imaging, embryos were imaged on a Zeiss Discovery V12 stereo microscope and images were captured with an AxioCam MRc digital camera. All images were collected at 20x in 12 Bit using an Achromat S 1.0x objective. Flurophores were excited using 488 (EGFP) and 554 (tdTomato) filters. Representative images of individual embryos are shown in Figure S2.

#### Mouse injections

Mouse embryos were injected using a digital injection system (Xenoworks, Sutter) and borosilicate filamented glass needles (Sutter, NC9955576) that were prepared in house. The microinjection system consists of a DMi8 microscope (Leica) equipped with micromanipulators (Sutter). Embryos were injected in glass bottomed dishes (MatTek, P35GC-1.5-14-C) in 25 μL drops of M2 covered in culture oil. First, 50 ng/μL of split fluorescent ALFA mRNA was injected into the zygote immediately after isolation, with 100 ng/μL of ALFAgrad mRNA injected into one cell of the two cell embryos 24 hours later. Embryos were fixed 12 h post degron injection. For fixation, the zona pellucida was first removed using acidified Tyrode solution (Sigma, T1788) and embryos were attached to a glass bottom dish (MatTek, P35GC-1.5-14-C). A solution of 4% PFA/PBS was added to embryos for 15 minutes at room temperature, which was followed by nuclear staining using 5 μg/ml DAPI (D1306, Thermo Fisher Scientific) for 15 minutes and imaging.

#### Mouse imaging

Mouse degron images were acquired using a Zeiss LMS 880 Airyscan inverted confocal microscope (Zeiss) using a C-Apochromat 40x/1.2 water objective with unidirectional line scanning at a format of 1476 x 1476 pixels, corresponding to an image size of 83.66 μm x 83.66 μm and 2.5x optical zoom. All images were collected at 16 Bit in sequential mode, and optical stacks were acquired at 1 μm spacing. Fluorophores were excited using the 405 laser line (DAPI) at 4.78% laser power, the 561 laser line (tdTomato) at 53.7% laser power and the 488 laser line (EGFP) at 47.1% laser power. Raw images were deconvolved using the Airyscan software. Representative slices of individual embryos are shown in Figure 2.

### Epitope tag knock-in

#### sgRNA design and synthesis

To target the N-terminus of Nanog and Vangl2, sgRNAs were designed to target as close to the start ATG codon as possible using CRISPRscan^59^. The sgRNA sequences were ordered from Synthego, resuspended to 500 ng/μL stocks in RNase-free H2O and stored at -80°C until use.

#### Nuclease test

A 5 μl mix containing 500 ng recombinant Engen Spy Cas9 NLS (NEB), 250 ng sgRNA and 300 mM KCl was incubated at 37°C for 5 minutes to form ribonucleoprotein complexes (RNPs). Embryos were injected at the 1-cell stage into the cell with the RNPs and grown to 48 hpf. Uninjected and injected embryos were lysed in a 1x PCR buffer and 1 μg/μL Proteinase K solution for 1 hour at 55°C followed by inactivation for 10 minutes at 95°C. 1 μL of this lysis was used in a PCR reaction that overlapped the target site listed below. The reactions were cleaned up using Zymo PCR Purification columns (Zymo) and Sanger sequenced. The resulting chromatograms were used for ICE analysis (Synthego) to determine cleavage efficiency. Each sgRNA was confirmed to have >80% cleavage efficiency across 3 independent F0 injected embryos. Targeting sgRNA sequence for *nanog* was TTTATCTAACGGCGAAATGG and for vangl2 was TGCGACTCGTTATCCATGTC. Genotyping for *nanog* was performed using the following primers, F: GTTGTAGGACAGAAAGAGCCGT, R: CACCTGGCAATATAAATCAGCA. Genotyping for *vangl2* was performed using the following primers, F: CCGCGCTCTCCAGTCCGTCA, R: CGAGAGCTGCGTGAGTGTGAA.

#### Tag knock-in design

Single stranded oligo donor nucleotides (ssODNs) were designed by inserting epitope sequences in frame with *nanog* and *vangl2* coding sequence directly proceeding the ATG start codon. Homology arms of 20 bp were appended to epitope tag sequences on the sense strand and ordered as PAGE-purified oligos (Sigma). Oligos were resuspended in RNase-free H2O at a concentration of 50 μM and stored at -20°C until use.

#### Tag knock-in injection

A 5 μL mix containing 2 μM donor oligo in RNase-free H_2_O was prepared. A second 5 μl mix containing 500 ng recombinant Engen Spy Cas9 NLS (NEB), 250 ng sgRNA and 300 mM KCl was incubated at 37°C for 5 minutes to form RNPs. Wildtype embryos were injected at the 1-cell stage twice: first, embryos were injected in the yolk with 1 nL of the donor oligo solution, and then second, embryos were re-injected into the cell with the RNPs. Uninjected, donor-only injected and dual injected embryos were grown to 48 hpf. 5 embryos per condition (uninjected, donor-only and dual injection) were lysed in a 1x PCR buffer and 1 μg/μL Proteinase K solution for 1 hour at 55°C followed by inactivation for 10 minutes at 95°C. 1 μL of this lysis was used in a PCR reaction that used *nanog* and *vangl2* genotyping primers with tag-specific primers, such that two junction PCRs were run for each sample. Clutches that showed positive junction PCR amplification were grown to adulthood for germline screening. Genotyping for ALFA integration was performed using the following primers, F: CACGTCTGGAGGAAGAGCTG, R: GTTCGGTCAGGCGACGAC.

Genotyping for VHH05 integration was performed using the following primers, F: TGGCAAGACAGATTAGCCCC, R: TTAGCCTCCTGATCAGCCTG. Genotyping for 127d01 integration was performed using the following primers, F: TCTGGAAGGGTGAGGATCCT, R: TCACCCTTCCAGAAATCTTCGA.

Donor ssODNs for *nanog* were the following:

**ALFA**: GAGTTTATCTAACGGCGAAATGCCATCACGTCTGGAGGAAGAGCTGCGTCGTCGCCTGACCGAACCTGCGGACTGGAAGATGCCA

**VHH05**: GAGTTTATCTAACGGCGAAATGCCGCAGGCTGATCAGGAGGCTAAAGAGCTGGCAAGACAGATTAGCCCCGCGGACTGGAAGATGCC A

**127d01**: GAGTTTATCTAACGGCGAAATGCCATCCTTCGAAGATTTCTGGAAGGGTGAGGATCCTGCGGACTGGAAGATGCCA

Donor ssODNs for *vangl2* were the following:

**ALFA**: CCCGCCCACTGGCCCCCGACATGCCATCACGTCTGGAGGAAGAG CTGCGTCGTCGCCTGACCGAACCTGATAACGAGTCGCAGTACTC

**VHH05**: CCCGCCCACTGGCCCCCGACATGCCGCAGGCTGATCAGGAGGCT AAAGAGCTGGCAAGACAGATTAGCCCCGATAACGAGTCGCAGTA CTC

**127d01**: CCCGCCCACTGGCCCCCGACATGCCATCCTTCGAAGATTTCTGGAAGGGTGAGGATCCTGATAACGAGTCGCAGTACTC

#### Line establishment

Adult F0 animals were incrossed to increase throughput of screening. Pools of 8 embryos were lysed in 50 μL 1x PCR buffer and 1 μg/μL Proteinase K solution for 1 hour at 55°C followed by inactivation for 10 minutes at 95°C. 12 pools of embryos per pair to maximize screening efficiency. F0 pairs that had both positive 5’ and 3’ were then outcrossed to wildtype fish, and 24 individual embryos from those crosses were re-screened by PCR. Embryos that were positive for both 5’ and 3’ junction PCRs were Sanger sequenced to assess tag integration. Those with precise integrations were grown to adulthood, genotyped for tag integration and incrossed to generate homozygous knock-in animals. Homozygous animals were further confirmed for precise tag integration by PCR using genotyping primers listed above. For all alleles, animals were maintained at homozyogosity for the knock-in allele and exhibited normal development.

### Endogenous ALFA-tagged protein imaging

Wildtype, ALFA-*nanog*^*KI/KI*^ and ALFA-*vangl2*^*KI/KI*^ were dechorionated in pronase and injected at the 1-cell stage with 25 pg membrane TagBFP2 (Addgene #55295) and 50 pg EGFP-NbALFA. Embryos were incubated at 28°C until they reached 30% epiboly (∼5 hpf) and mounted in 0.8% low melt agarose (GPG/LMP AmericanBio, AB00981-00050) in system water against a no. 1.5 cover slip. Embryos were imaged on a temperature controlled (28°C) upright Zeiss LSM 980 confocal microscope with an Airyscan 2 detector and a Plan-Apochromat 20x/0.8 M2 objective with 2.50 second scan speed, bidirectional line scanning at a format of 4060 x 4060 pixels and 1.4x optical zoom. All images were collected at 16 Bit, and optical stacks were acquired at 0.3 μm spacing. Fluorophores were excited using the 408 laser line (mTagBFP2) at 2% laser power and 488 laser line (EGFP) at 1% laser power. Raw images were deconvolved using the Airyscan software. Individual representative slices of the enveloping layer are shown in Figure 3.

### Endogenous ALFA-tagged protein depletion

Wildtype, ALFA-*nanog*^*KI/KI*^ and ALFA-*vangl2*^*KI/KI*^ were dechorionated in pronase and injected at the 1-cell stage with 50 pg ALFAgrad mRNA. For ALFA-*nanog*^*KI/KI*^, embryos were incubated at 28°C and scored at 6 hpf for *nanog* loss-of-function phenotypes (gastrulation failure). For ALFA-*vangl2*^*KI/KI*^, embryos were incubated at 28°C and scored at 24 hpf for *vangl2* loss-of-function phenotypes (shortened body axis).

### Western blot

#### Comparison to overexpression

Wildtype embryos were injected with 25 and 50 pg of ALFA-*nanog* mRNA at the 1-cell stage and incubated at 28°C until they reached 50% epiboly. 25 embryos from uninjected, 25 pg injected, 50 pg injected and stage-matched ALFA-*nanog*^*KI/KI*^ were batch deyolked in deyolking buffer (55 mM NaCl, 1.8 mM KCl, 1.25 mM NaHCO_3_) and snap frozen in liquid nitrogen. The embryos were boiled in sample buffer (1x NuPAGE LDS Sample Buffer, 1x NuPAGE Sample Reducing Agent (ThermoFisher)) for 10 min. Samples were resolved on a 4-12% NuPAGE Bis-Tris gel in NuPAGE MOPS Running Buffer (Thermo Fisher) and transferred to a nitrocellulose membrane with the iBlot 2 Gel Transfer Device (Thermo Fisher). The membrane was blocked in 5% milk / PBS with 0.1% Tween-20 (PBS-Tw) and then cut at ∼25 kDa. The membranes were incubated with primary antibody solution (1:1000 anti-ALFA antibody (NanoTag Biotechnologies) for the top (>25 kDa) membrane, 1:5000 anti-H3 antibody (Abcam, ab1791) for the bottom (<25 kDa) membrane) prepared in blocking solution), and then incubated with secondary antibody solution (1:10000 of horseradish peroxidase-conjugated anti-rabbit antibody (Abcam)). The Super-Signal West Pico PLUS Chemiluminescent Substrate (Thermo Fisher) was used for protein detection. Fiji software was used for densitometry to assess levels of ALFA-tagged Nanog in different samples normalized to histone H3 loading control.

#### Time course

For each timepoint, 15 ALFA-*nanog*^*KI/KI*^ embryos were dechorionated by hand in Ringer’s solution (116 mM NaCl, 2.9 mM KCl, 1.8 mM CaCl_2_, 5.0 mM HEPES, pH 7.2) and rinsed in PBS. The embryos were boiled in sample buffer (1x NuPAGE LDS Sample Buffer, 1x NuPAGE Sample Reducing Agent (ThermoFisher)) for 10 min. The samples were resolved on a 4-12% NuPAGE Bis-Tris gel in NuPAGE MOPS Running Buffer (Thermo Fisher) and transferred to a nitrocellulose membrane with the iBlot 2 Gel Transfer Device (Thermo Fisher). The blot was processed the same as above.

### Chromatin Immunoprecipitation Sequencing (ChIP-seq)

ChIP-seq was performed as described^29,60^. Briefly, embryos from ALFA-*nanog*^*KI/KI*^ females crossed with ALFA-*nanog*^*KI/KI*^ males were dechorionated at the 1-cell stage. At 4 hpf, 700 embryos were fixed with 1.9% paraformaldehyde for 15 minutes at room temperature, quenched with 0.125 M glycine for 5 minutes, washed 3 times with cold PBS and snap-frozen with liquid nitrogen). The ALFA-*nanog*^*KI/KI*^ embryos were homogenized and lysed for 15 minutes in cell lysis buffer (10 mM Tris–HCl pH 7.5, 10 mM NaCl, 0.5% IGEPAL, protease inhibitors) on ice. Nuclei were precipitated by centrifugation for 5 minutes at 3500 rpm at 4°C. Nuclei were then lysed with nuclear lysis buffer for 10 minutes (50 mM Tris–HCl pH 7.5, 10 mM EDTA, 1% SDS, protease inhibitors) on ice, diluted with IP dilution buffer (16.7 mM Tris–HCl pH 7.5, 167 mM NaCl, 1.2 mM EDTA, 0.01% SDS, protease inhibitors), and sonicated (15 cycles of sonication with 30 seconds ON and 30 seconds OFF, 15 minutes in ice, and another 15 cycles of sonication; Bioruptor Pico sonication device, Diagenode). 8 μl of 10% Triton X-100 was added per 100 μl of sonicated chromatin to the chromatin suspension, which was then centrifuged for 10 minutes at 14,000 rpm at 4°C. 5% of the supernatant was taken as input and stored at -80°C until use. 25 μl of Protein G Dynabeads were washed with 0.5% BSA/PBS, incubated with 4 ug of ALFA antibodies from Nanotag Biotechnologies (4 μg of N1583 + 2 μg of N1581) overnight at 4°C, washed three times with cold 0.5% BSA/PBS, and added to the chromatin supernatant to incubate overnight at 4°C. The beads were washed five times with cold RIPA wash buffer (50 mM HEPES pH 7.6, 1 mM EDTA, 0.7% DOC, 1% Igepal, 0.5 M LiCl), two times with TBS (Tris-buffered saline; 50 mM Tris pH 7.5, 150 mM NaCl) before eluted with elution buffer at 65°C for 15 minutes (50 mM NaHCO_3_, 1% SDS). Both the input and ChIP samples were purified for sequencing: reverse-crosslinking at 65°C overnight, RNase A (0.33 μg/μl) treatment at 37°C for 2 hours, Proteinase K (0.2 μg/μl) treatment at 55°C for 2 hours, and purification with ChIP DNA Clean & Concentrator (D5205, Zymo Research). Library preparation (Illumina TruSeq protocol) and sequencing (Illumina NovaSeq 6000 System, pair-end) were performed by the Yale Center for Genome Analysis.

### Immunofluorescence of ALFA-Nanog

ALFA-*nanog*^*KI/KI*^ were dechorionated in pronase at the 1-cell stage and grown at grown at 28°C until they reached sphere stage (∼4 hpf). Embryos were fixed in 4% PFA in PBS overnight at 4°C, followed by 1 wash in 1xPBS with 0.1% Triton-X (PBS-Tx) for 10 minutes at room temperature. Embryos were blocked in 10% normal goat serum (NGS) (Thermo Fisher Scientific, 50062Z) for 2 hours with gentle rotation and incubated with primary antibody solution (1:1000 anti-ALFA antibody (NanoTag Biotechnologies) prepared in 10% NGS overnight at 4°C. Embryos were washed 3 times for 30 minutes in PBS-Tx, followed by incubation with secondary antibody solution (1:1000 Goat anti-Rabbit Alexa Fluor 488 (Thermo Fisher Scientific, A-11008) and 1:1000 DAPI) with gentle rotation for 1 hour at room temperature protected from light. Embryos were washed 3 times for 30 minutes in PBS-Tx and mounted in 0.8% low-melt agarose on glass-bottom dishes (MatTek). Embryos were imaged on an upright Zeiss LSM 980 confocal microscope with an Airyscan 2 detector and a Plan-Apochromat 20x/0.8 M2 objective with bidirectional line scanning at a format of 4084 x 4084 pixels and 1.3x optical zoom. All images were collected at 16 Bit, and optical stacks were acquired at 3.08 μm spacing. Fluorophores were excited using the 405 laser line (DAPI) at 0.5% laser power and 488 laser line (EGFP) at 7.5% laser power. Raw images were deconvolved using the Airyscan software. Representative orthogonal projections of individual embryos are shown in Figure S3.

### Live image acquisition

For live imaging of Nanog, ALFA-*nanog*^*KI/KI*^ embryos were dechorionated with pronase and injected with 75 pg of the EGFP-NbALFA GEAR in RNase-free H_2_O at the 1-cell stage. The exogenous ALFA-*nanog* mRNA was injected at 25 pg together with 75 pg of the EGFP-NbALFA GEAR and the exogenous *nanog*-mEmerald^42^ reporter was injected at 25 pg. Injected embryos were grown at 28°C and mounted in 0.8% low melt agarose (GPG/LMP AmericanBio, AB00981-00050) in system water at the 4-cell stage against a no. 1.5 cover slip. Temperature controlled live imaging (28°C) was performed using an upright Zeiss LSM 980 confocal microscope with an Airyscan 2 detector and a LD LCI Plan-Apochromat 40x/1.2 Imm Corr DIC M2 objective with water immersion. Time series were acquired starting at the 8-cell stage and confocal stacks were readjusted during mitosis of every cleavage cycle. In detail, images were obtained at 16 Bit with 2x line averaging, bidirectional scanning, LSM scan speed 8 (pixel dwell time 0.37 μs), 2x optical zoom and using an image size of 2124×2124 pixel corresponding to an image size of 105.47×105.47 μm. EGFP was excited using the 488 nm laser line at 2.8% laser power. Three-dimensional optical sections were acquired at 1 μm distance, a final depth of 31 μm and a final temporal resolution of 32 seconds per time frame.

Imaging of Nanog in conjunction with elongating Pol II (Pol II pSer2 mintbody^42^) was carried out using the microscope setup described above with the following alterations. Injections of 75 pg mScarlet-i3-NbALFA GEAR and 25 pg pSer2-EGFP mintbody into ALFA-*nanog*^*KI/KI*^ embryos were performed after dechorionation at the 1-cell stage. In total, 61 planes were acquired with 1 μm spacing (total 60 μm), optical zoom of 1.5, a z-stack acquisition time of ∼147 sec and a format of 2848×2848 pixels corresponding to an image size of 140.83×140.83 μm. mScarlet-i3 was excited by the 561 nm laser line at 1.1% laser power and the mintbody by the 488nm laser line at 2.4%.

Maximum intensity projections of 3D stacks are shown in the result sections. Embryos were imaged for 120-180 min and included the beginning of the 1k cell stage. During image analysis all datasets were adjusted in time to account for slight temperature differences during imaging that can alter the speed of development. Therefore, each cell cycle was aligned to start with the completion of telophase and ended with chromatin decondensation.

## Quantification and Statistical Analysis

### Degron analysis

#### Zebrafish reporters at 10 hpf

3D image analysis was performed in the IMARIS software (Bitplane, Oxford Instruments, Concord MA; Version: 10.0). Nuclei were identified using the ‘spot’ function and a constant size of 5 μm in xy and a z-axis point spread function of 8μm. The membrane fluorescence of a cell was quantified by sampling the membrane circumference in multiple regions using the ‘spots’ function. Smaller spots with a size of 2 μm xy and 4 μm in the z-dimension were added based on the tdTomato staining. To determine the background fluorescence in both channels, sets of ‘spots’ of the same volume as the nuclei and membrane spots were generated for background correction. Statistics for all spot objects were exported and membrane spots were linked to their closest nucleus in 3D using a custom python script^61^. Median background fluorescence was subtracted from each channel and the median nuclear sum fluorescence divided by the median membrane sum fluorescence for each cell. Data for all individual replicates can be found plotted in Figure S2 H-J. In Figure 2, the median nuclear to membrane fluorescence ratio of each embryo was plotted and the resulting mean of the control embryos was set to 1 and all other values were adjusted accordingly.

#### Zebrafish reporters at 24 hpf

The total GFP or tdTomato intensities (I_total_) were measured using a 0.1 x 0.2 inch rectangle in the trunk of the embryo above the yolk extension. The average background intensity (I_background_) was measured for each channel and embryo using a 0.1 x 0.2 inch rectangle adjacent to the embryo. The average intensity of the (I_average_) was measured by subtracting I_background_ from I_total_. The fold decrease in GFP expression was calculated as a ratio of the GFP intensity in degron-injected and uninjected embryos, divided by the ratio of tdTomato intensity in degron-injected and uninjected embryos to normalize for differences due to injection.

#### Mouse

Two-cell mouse embryos were used for quantification purposes where one cell served as a direct control to the second cell that was injected with the degrader. In the IMARIS image analysis software nuclei were segmented using the ‘spots’ module based on the DAPI signal and an diameter of 10 μm. Membrane fluorescence was sampled for each cell by adding 2 μm (xy) x 4 μm (z) spots based on the tdTomato fluorescence. The cell membrane portion shared between the two cells was excluded from analysis, leaving only cell membrane portions unique to the individual cells.

### ALFA-Nanog immunofluorescence quantification

Nuclear Nanog concentration was determined based on Ab-ALFA immunofluorescence staining. Nuclei were segmented in the IMARIS software using the ‘surface’ module and based on the DAPI channel. The sum fluorescence in the Ab-ALFA channel of each nucleus was adjusted for the respective nuclear volume and background corrected to account for background variation between experiments. The median background fluorescence for each embryo was determined by placing 3×3 μm spheres using the ‘spot’ function randomly within the embryo (excluding nuclear regions).

### ChIP-seq analysis

The ChIP-seq data were managed with LabxDB seq^62^. To validate our endogenous ALFA-Nanog ChIP-seq (4 hpf), we also analyzed the 4.5 hpf Myc-Nanog ChIP-seq that was done by overexpressing Myc-tagged Nanog in wildtype embryos^41^. Raw reads were mapped to the zebrafish GRCz11 genome sequence^63^ with LabxPipe (https://github.com/vejnar/LabxPipe) using Bowtie2^64^ (option: --no-unal, --no-discordant, --no-mixed, version: 2.5.1) after adapter-trimming (using ReadKnead, https://github.com/vejnar/ReadKnead). Reads were filtered (deduplicated, and only uniquely mapped reads (MAPQ ≥ 30) were kept) using SAMtools^65^ before any downstream analysis. Reads from the single-end Myc-Nanog ChIP-seq^41^ were extended to 200 nt for all downstream analyses. Genomic tracks were created using the LabxPipe trackhub option, which uses GeneAbacus (https://github.com/vejnar/GeneAbacus) to compute fragment coverage (normalized to the total fragments per million fragments). To plot the correlation between the 4 hpf ALFA-Nanog ChIP-seq and the 4.5 hpf Myc-Nanog ChIP-seq, the average ChIP-seq signal were calculated across each 5-kb window of the zebrafish genome; Pearson correlation was then calculated on all genomic windows. Narrow peaks were called with input data as controls from the filtered BAM files (significance cut-off: q = 0.05; options specific for the single-end Myc-Nanog ChIP-seq: “-f BAM --nomodel --extsize 200”; options specific for ALFA-Nanog ChIP-seq: “-f BAMPE”) using MACS3^66^. Single nucleotide resolution heatmaps, centered at the peak centers, were created with deepTools^67^.

### Live imaging analysis

#### Nanog foci quantification

Nanog foci segmentation of live imaging data was performed using the IMARIS software. Foci were defined to have an x-y diameter of 0.5 μm and a z-length of 1 μm and identified using the ‘spot’ function. Spots were thresholded based on their fluorescence intensity over background. To account for injection variability and fluorescence heterogeneity between experiments, random background fluorescence measurements were taken in the embryo in the last frame before chromatin condensation of each cell cycle. The median background fluorescence of an embryo was subtracted from the sum fluorescent values of Nanog foci. Resulting fluorescence values below a cutoff (65000 (au) ∼ lowest 6% of dynamic range) were discarded.

#### Nanog foci distance to active transcription sites

Analysis was performed in the IMARIS software. First nuclei were segmented based on their nuclear fluorescence intensity using the ‘surface’ module. Each nucleus was then “immobilized” through drift correction in x/y/z dimensions based on its centroid. Nanog foci and transcription foci were segmented using the ‘spots’ module and their x/y/z centroid position was extracted. The shortest distance in 3D between Nanog foci and their respective closest transcription spot was computed.

### Statistical analysis

Statistical comparisons were performed using two-tailed Student’s t tests, Mann-Whitney tests and one-way ANOVA with Dunnett’s multiple comparisons tests in GraphPad Prism 10. ChIP-seq analysis and Pearson Correlation was performed using Python. Statistical test and sample sizes can be found in Figure legends. Statistical significance was assumed by p<0.05. Individual p values are indicated in Figure legends.

